# Deep sequencing of 3 cancer cell lines on 2 sequencing platforms

**DOI:** 10.1101/623702

**Authors:** Kanika Arora, Minita Shah, Molly Johnson, Rashesh Sanghvi, Jennifer Shelton, Kshithija Nagulapalli, Dayna M. Oschwald, Michael C. Zody, Soren Germer, Vaidehi Jobanputra, Jade Carter, Nicolas Robine

## Abstract

To test the performance of a new sequencing platform, develop an updated somatic calling pipeline and establish a reference for future benchmarking experiments, we sequenced 3 common cancer cell lines along with their matched normal cell lines to great sequencing depths (up to 278X coverage) on both Illumina HiSeqX and NovaSeq sequencing instruments. Somatic calling was generally consistent between the two platforms despite minor differences at the read level. We designed and implemented a novel pipeline for the analysis of tumor-normal samples, using multiple variant callers. We show that coupled with a high-confidence filtering strategy, it improves the accuracy of somatic calls. We also demonstrate the utility of the dataset by creating an artificial purity ladder to evaluate the somatic pipeline and benchmark methods for estimating purity and ploidy from tumor-normal pairs. The data and results of the pipeline are made accessible to the cancer genomics community.

## Introduction

The field of cancer genomics has exploded with the development of high-throughput sequencing, largely driven by Illumina’s short read sequencing technology. Thousands of tumors have been sequenced in the last decade, with strategies varying from variant hotspot panels [1], cancer gene panels [2], whole-exome (such as used in The Cancer Genome Atlas project [3]) or whole-genome sequencing (WGS) [4]. In 2014, Illumina introduced the HiSeq X Ten (HiSeqX) as their main sequencing instrument dedicated to human whole-genome sequencing. In 2017, they released the NovaSeq 6000 Sequencing System (NovaSeq), which is currently the latest generation of Illumina sequencing instruments. The primary difference between these platforms is the adoption of 2-channel Sequencing-by-Synthesis in the NovaSeq where clusters detected in the red wavelength filter correspond to a C nucleotide, clusters detected in the green wavelength filter correspond to a T, clusters detected by both colors correspond to A, and unlabeled clusters are G bases. For the NovaSeq, Illumina introduced a new base calling algorithm and method for estimating Quality scores, with 4 quality bins (as opposed to 8 bins for HiSeqX).

With the introduction of any new sequencing technology, it is important to investigate the error profiles and biases of the technology, and to understand the subsequent impact of those on downstream analyses. This is especially important for cancer data analysis where varying tumor purity and intra-tumor heterogeneity make distinguishing low frequency somatic variants from sequencing noise challenging. Here, we have created a whole genome reference dataset of 3 matched tumor-normal cell lines sequenced deeply on both HiSeqX and NovaSeq, employed it to calibrate our somatic pipeline, and released it to the genomics community. The 3 cancer cell lines selected are common and represent the range of mutations profiles that a somatic pipeline is expected to identify correctly. COLO-829 was derived from a metastatic melanoma male patient and presents a pseudo-tetraploid karyotype [5] [6]. It was the first cancer genome to be comprehensively characterized by whole-genome sequencing [7], has previously been characterized as hypermutated and was used to establish a reference for benchmarking somatic mutation pipelines [8]. HCC-1143 and HCC-1187 were isolated from patients with ductal carcinoma breast cancer [9]. HCC-1143 is near tetraploid [10] and heavily rearranged, and its matched normal cell line HCC-1143BL has a chromosome 2 amplification. HCC-1187 is hypotriploid [11].

## Methods

### Cell culture and DNA isolation

The cancer cell lines (COLO-829 ATCC® CRL-1974™, COLO 829BL ATCC® CRL-1980™, HCC-1143 ATCC® CRL-2321™, HCC1143 BL ATCC® CRL-2362™, HCC-1187 ATCC®CRL-2322™ and HCC-1187BL ATCC® CRL-2323™) were obtained from ATCC^2^. The cell lines were cultured using the recommendations from ATCC. Cultured cells were split into two aliquots for metaphase chromosome preparation and karyotype analysis. Representative images and karyotypes are reported in Supplemental Figure S1. The number of passages is indicated in Supplemental Table 1.

### Library preparation and sequencing

Libraries were prepared using the TruSeq DNA PCR-free Library Preparation Kit (Illumina) with 1μg DNA input following Illumina’s recommended protocol^3^, with minor modifications as described. Intact genomic DNA was concentration normalized and sheared using the Covaris LE220 sonicator to a target size of 450bp. After cleanup and end-repair, an additional double-sided bead-based size selection was added to produce sequencing libraries with highly consistent insert sizes. This was followed by A-tailing, ligation of Illumina DNA Adapter Plate (DAP) adapters and two post-ligation bead-based library cleanups. These stringent cleanups resulted in a narrow library size distribution and the removal of remaining unligated adapters. Final libraries were run on the Fragment Analyzer to assess their size distribution and quantified by qPCR with adapter specific primers (Kapa Biosystems). The libraries were pooled together based on expected final coverage and sequenced across multiple flow cell lanes to reduce impact of lane-to-lane variations in yield. Whole genome sequencing was performed on the NovaSeq (NSCS 1.3.1; RTA v3.3.3) and the HiSeqX (HCS HD 3.5.0.7; RTA v2.7.7) at 2×150 bp read length, using v1 S2/S4 300-cycle and SBS v3 reagents, respectively.

### Pre-processing

The sequencing data for all the cell lines was demultiplexed using bcl2fastq (Illumina) v2.20.0.422. FASTQ files were then processed through NYGC’s high-performance computing cluster using the NYGC automated pipeline (Figure 1). Sequencing reads were aligned to the GRCh38 reference genome (1000 Genomes version) using BWA-MEM (v0.7.15) [12]. NYGC’s ShortAlignmentMarking (v2.1)^4^ was used to mark short reads as unaligned. This tool is intended to remove spurious alignments resulting from contamination (e.g. saliva sample bacterial content) or from too aggressive alignments of short reads the size of BWA-MEM’s 19bp minimum seed length. These spurious alignments result in pileups in certain locations of the genome and can lead to erroneous variant calling.

**Figure 1:**
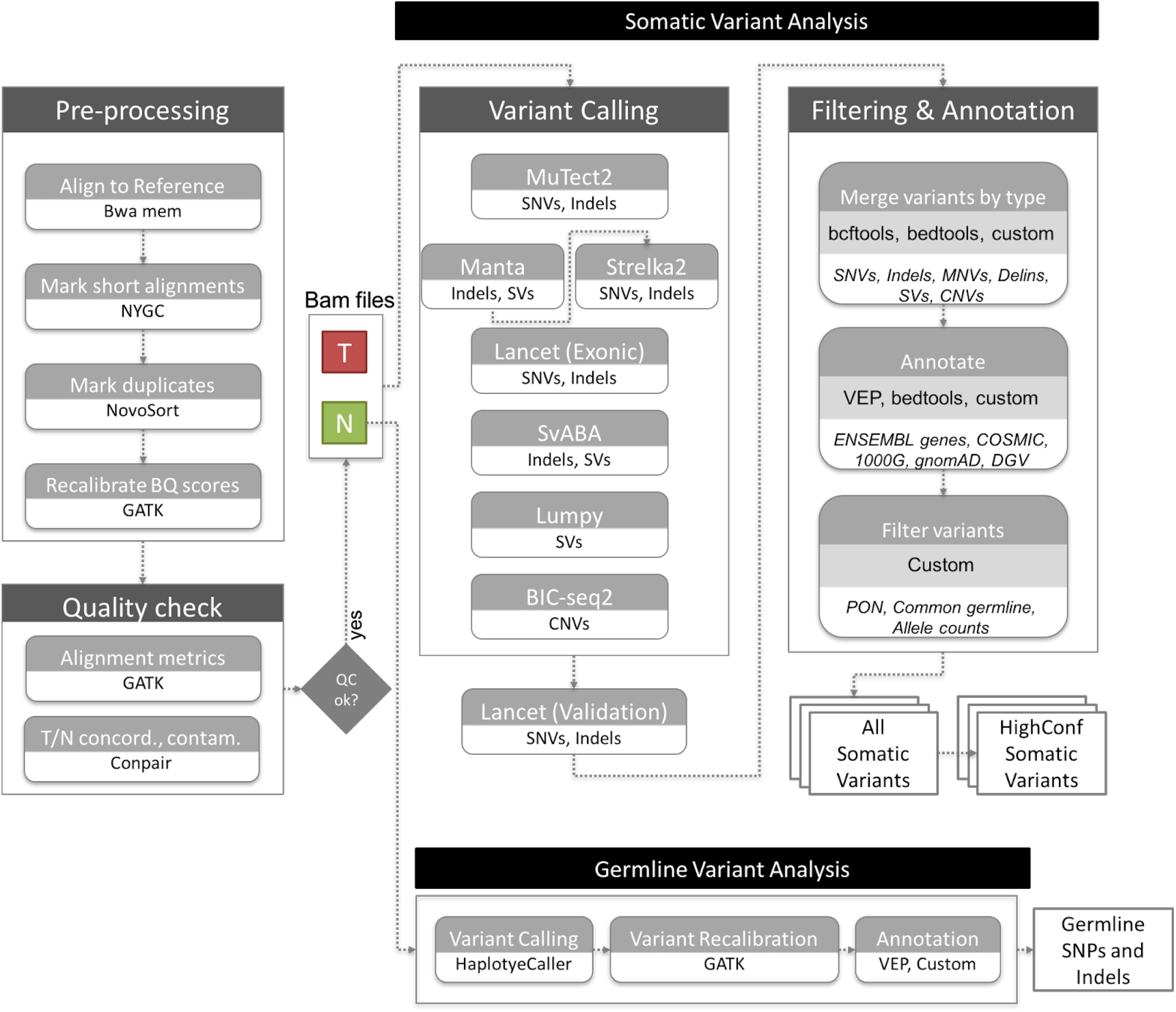
NYGC Somatic Pipeline for tumor-normal whole-genome sequencing samples.

GATK (v4.1.0) [13] FixMateInformation was run to verify and fix mate-pair information, followed by Novosort (v1.03.01) markDuplicates to merge individual lane BAM files into a single BAM file per sample. Duplicates were then sorted and marked, and GATK’s base quality score recalibration (BQSR) was performed. The final result of the pre-processing pipeline was a coordinate sorted BAM file for each sample.

Once preprocessing was complete, we computed a number of alignment quality metrics such as average coverage, %mapped reads and %duplicate reads using GATK (v4.1.0) and an autocorrelation metric (adapted for WGS from[14]) to check for unevenness of coverage. We also ran Conpair[15], a tool developed at NYGC to check the genetic concordance between the normal and the tumor sample and to estimate any inter-individual contamination in the samples.

### Variant calling and annotation

The tumor and normal bam files were processed through NYGC’s variant calling pipeline which consists of MuTect2 (GATK v4.0.5.1) [16], Strelka2 (v2.9.3) [17] and Lancet (v1.0.7) [18] for calling Single Nucleotide Variants (SNVs) and short Insertion-or-Deletion (Indels), SvABA (v0.2.1) [19] for calling Indels and Structural variants (SVs), Manta (v1.4.0) [20] and Lumpy (v0.2.13) [21] for calling SVs and BIC-Seq2 (v0.2.6) [22] for calling Copy-number variants (CNVs). Manta also outputs a candidate set of Indels which is provided as input to Strelka2 (following the developers recommendation, as it improves Strelka2’s sensitivity for calling indels >20nt). Due to its computing requirements, in this pipeline Lancet is only run on the exonic part of the genome. It is also run on the +/-250nt regions around non-exonic variants that are called by only one of the other callers, to add confidence to such variants. Small SVs called by Manta are also used to add confidence to the indel calls.

Next, the calls were merged by variant type (SNVs, Multi Nucleotide Variants (MNVs), Indels and SVs). MuTect2 and Lancet call MNVs, however Strelka2 does not and it also does not provide any phasing information. So to merge such variants across callers, we first split the MNVs called by MuTect2 and Lancet to SNVs, and then merged the SNV callsets across the different callers. If the caller support for each SNV in a MNV is the same, we merged them back to MNVs. Otherwise those were represented as individual SNVs in the final callset. The SVs were converted to bedpe format, all SVs below 500bp were excluded and the rest were merged across callers using bedtools pairtopair (slop of 300bp, same strand orientation, and 50% reciprocal overlap). For CNVs, segments with log2 > 0.2 were categorized as amplifications, and segments with log2 < −0.235 were categorized as deletions (corresponding to a single copy change at 30% purity in a diploid genome, or a 15% Variant Allele Fraction). The resulting variants were annotated with Ensembl as well as databases such as COSMIC (v86) [23], 1000Genomes (Phase3)[24], gnomAD (r2.0.1) [25], dbSNP (v150) [26] and Database of Genomic Variants (DGV) [27] using Variant Effect Predictor (v93.2) [28] for SNVs and Indels, and bedtools [29] for SVs and CNVs.

### Somatic variant filtering

#### Panel Of Normals

The Panel Of Normals (PON) filtering removes recurrent technical artifacts from the somatic variant callset [16]. The Panel of Normals for SNVs, indels and SVs was created with whole-genome sequencing data from normal samples from 242 unrelated individuals. Of these, sequencing data for 148 individuals was obtained from the Illumina Polaris project^5^ which was sequenced on the HiSeqX platform with PCR-free sample preparation. The remaining samples were sequenced by the NYGC. Of these, 73 individuals were sequenced on HiSeqX, 11 on NovaSeq, and 10 were sequenced on both.

We ran MuTect2 in artifact detection mode and Lumpy in single sample mode on these samples. For SNVs and indels, we created a PON list file with sites that were seen in two or more individuals and we used this list to filter the somatic variants in the merged SNV and indel files.

For SVs, we used SURVIVOR (v1.0.3) [30] to merge Lumpy calls. Variants were merged if they were of the same type, had the same strand orientation, and were within 300bp of each other (maximum distance). We did not specify a minimum size. After merging SVs, we used these calls as a PON list. To filter our somatic SV callset, we merged our PON list with our callset using bedtools pairtopair (slop of 300bp, same strand orientation, and 50% reciprocal overlap), and filtered those SVs found in two or more individuals in our PON.

#### Common germline variants

In addition to the PON filtering, we removed SNVs and Indels that had minor allele frequency (MAF) of 1% or higher in either 1000Genomes (phase 3) or gnomAD (r2.0.1) [25], and SVs that overlapped DGV and 1000Genomes (phase3). CNVs were annotated with DGV and 1000 Genomes but not filtered.

#### Allele counts

Since our variant callsets were generated by merging calls across callers, and each of them reported different allele counts, we reported final chosen allele counts for SNVs and indels. For SNVs, and for indels less than 10nt in length, these were computed as the number of unique read-pairs supporting each allele using the pileup method, with minimum mapping quality and base quality thresholds of 10 each.

For larger indels and complex events, we chose the final allele counts reported by the individual callers Strelka2, MuTect2, Lancet, in that order. For indels larger than 10nt that were only called by SvABA, we did not report final allele counts and allele frequencies because SvABA does not report the reference allele count, making it difficult to estimate the variant allele frequency.

We then used these final chosen allele counts and frequencies to filter the somatic callset. Specifically, we filtered any variant for which the variant allele frequency (VAF) in the tumor sample was less than 0.0001, or if the VAF in the normal sample was greater than 0.2, or if the depth at the position was less than 2 in either the tumor sample or the normal sample. We also filtered variants for which the VAF in normal sample was greater than the VAF in tumor sample. Variants that passed all of the above-mentioned filters were included in our final somatic callset (hereby referred to as AllSomatic).

#### High-confidence variants

For SNVs, indels and SVs, we also annotated a subset of the somatic callset as high confidence. For SNVs and indels, high confidence calls were defined as those that were either called by two or more variant callers, or called by one caller and also seen in the Lancet validation calls or in the Manta SV calls.

For structural variants, high confidence calls were taken from the somatic callset if they met the following criteria: a SV was called by 2 or more variant callers, or called by Manta or Lumpy with either additional support from nearby CNV changepoint or split-read support from SplazerS [31], an independent tool used to calculate the number of split-reads supporting SV breakpoints. An SV was considered supported by SplazerS if it found at least 3 split-reads in the tumor only. Nearby CNV changepoints were determined by overlapping BIC-Seq2 calls with the SV callset using bedtools closest. An SV was considered to be supported by a CNV changepoint if the breakpoint of the CNV was within 1000bp of an SV breakpoint.

##### Germline variant analysis

We also called germline SNPs and indels using GATK HaplotypeCaller (v3.5), which generated a single-sample GVCF. We then ran GATK’s GenotypeGVCF to perform single sample genotype refinement and output a VCF, followed by variant quality score recalibration (VQSR) for variant filtering (at tranche 99.6%). Next, we ran Variant Effect Predictor (v93.2) to annotate the variants with Ensembl as well as databases such as COSMIC (v86), 1000Genomes (Phase3), gnomAD (r2.0.1), dbSNP (v150), ClinVar (201805), Polyphen2 (v2.2.2) and SIFT (v5.2.2).

### Mismatch analysis

The whole genome data from both platforms was downsampled to 8X coverage for all samples, and the number of single nucleotide mismatches to the reference in each sample was computed in order to evaluate the technical error profiles for each sequencing platform. Duplicate, unmapped, supplementary and vendor/platform QC-failed reads were excluded from the calculations. Mismatches were represented with respect to the read strand. For downstream analyses, mapping quality ≥10 and base quality ≥ 10 cut-offs were applied. The mismatch types were classified by the trinucleotide context in which they occur in the read strand. Mismatches were also summarized for the 6 reduced categories: C>A, C>G, C>T, T>A, T>C, T>G, by reverse complementing the other categories.

#### PCAWG Sanger pipeline

The CWL workflow definition file and the pre-compiled resource files were downloaded according to the instructions^6^. The pipeline was run using cwltool (v1.0.52) and only supports the GRCh37 reference.

#### Comparison to COLO-829 reference dataset

We downloaded the SNVs and Indels VCF file from EGA (Data accession ID: EGAD00001002142). This reference dataset was only provided for GRCh37. We therefore ran our pipeline and the PCAWG Sanger pipeline on the 80X/40X COLO-829/COLO-829BL HiSeqX and NovaSeq data aligned to GRCh37 with decoys reference (from GATK bundle for b37) and compared the somatic SNVs and Indels called on our data to the PASS-filtered variants in the Craig et al. VCF file. MNVs called by our pipeline were converted to SNVs for this comparison. CNV gene-level information was taken from Supplementary Table 2 of Craig et al. and compared to the CNV calls from our pipeline and the PCAWG Sanger pipeline after converting the data to gene-level results.

#### Data downsampling and purity ladder generation

Since the samples were sequenced to different depths between the two platforms, for the HiSeqX vs NovaSeq comparisons, we downsampled the tumor samples to 80X and the normal samples to 40X for all inter-platform variant comparisons. For this we used samtools view to randomly subsample the reads from the high coverage data.

For within platform, intra-run comparisons, we split the high coverage NovaSeq data for COLO-829, COLO-829BL, HCC-1143 and HCC-1143BL by readgroups using samtools split. Each readgroup corresponded to a different lane on the sequencer for that sample. Different sets of readgroups were used to create two replicates (Rep1 and Rep2) for each sample, thereby ensuring that the two replicate samples consisted of mutually exclusive set of reads. The tumor samples were then downsampled to 80X and normal samples to 40X.

To simulate low tumor purity samples, we downsampled the Rep1 NovaSeq data for the tumors, COLO-829 and HCC-1143, to 10X, 20X, 30X, 40X, 50X, 60X and 70X, and mixed that with data from their matched normal samples at 70X, 60X, 50X, 40X, 30X, 20X and 10X respectively. These mixed-in datasets were then analyzed against the 40X Rep1 matched normal sample data. Again, we used different readgroups for the mix-in than those that went into the Rep1 normal sample data.

To estimate the purity of these mixed-in samples, we had to take into consideration the average ploidy of the tumor. We used the ploidy that was estimated by CELLULOID [32] for the 100% purity samples, since they seemed to be very accurate upon manual review.

The tumor purity of the mixed-in samples was estimated using the following equation:

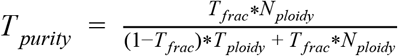

Where,

*N*_*ploidy*_ = Average ploidy of the normal sample, which we assumed to be 2.

*T*_*ploidy*_ = Average ploidy of the tumor sample, for which we used CELLULOID’s estimate of ploidy on the 100% purity data.

*T*_*frac*_ = Fraction of the reads in the mixed-in sample that came from the tumor. (For a 10X tumor+70X normal mix-in sample, this was 10/(10+70) = 0.125).

#### Purity Ladder Precision/Recall

SNVs and Indels called on the purity ladder samples were compared to the variants called in the high coverage data. True positives (TP) were considered to be variants that were also seen in the high confidence callset of the high coverage data, whereas false positives (FP) were considered to be variants that were called in the purity ladder sample but not seen in the AllSomatic callset of the high coverage data. Those variants that were called in the high confidence callset of the high coverage but not called in the purity ladder sample callset were classified as False Negatives (FN). Variants that were in the AllSomatic callset of the high coverage data but not in the HighConfidence callset were ignored for this analysis because they could not be confidently assigned as true variants.

For CNVs, events called in low purity samples were compared at the base level to the CNVs called in the high coverage 100% purity data. If a deletion or amplification was found in the low purity cell line, but not in the high coverage 100% purity cell line, this was classified as a FP. If a deletion/amplification was not found in the low purity cell line, but called in the high coverage 100% purity cell line, this was classified as a FN. TP were considered to be any deletions or amplifications that were found at the same position in both the low purity cell line and high coverage 100% purity cell line.

Precision, recall and F1 scores were calculated as:

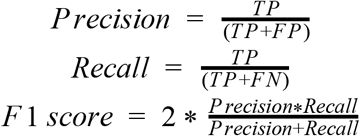

Purity-ploidy adjustment of CNV log2 values:

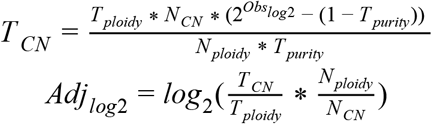

Where,

*T*_*CN*_= Absolute copy number for that segment in the tumor sample

*N*_*CN*_= Absolute copy number for that segment in the normal sample (2 for autosomes, 1 for sex chromosomes in male, 2 for X chromosome in female)

*N*_*ploidy*_= Average ploidy of the normal sample (which we assumed to be 2)

*T*_*ploidy*_= Average ploidy of the tumor sample

*T*_*purity*_= Tumor purity

*Obs*_*log2*_= Observed log2 value for the segment (from BIC-seq2)

*Adj*_*log2*_= Adjusted log2 value.

#### Purity-ploidy estimation using CELLULOID and HATCHet

For purity and ploidy estimation, we used CELLULOID (v0.11) and HATCHet^7^. CELLULOID was run in single-clone mode with default parameters and segment-based optimization. HATCHet was also run with default parameters.

## Results

### Read level comparison

We sequenced between 2 and 6.3 Billion reads for the 3 tumor cell lines, and between 1 and 4 Billion reads for the normal cell lines (Supplemental Table 2). Before applying our alignment and variant calling pipeline to the samples, we observed a few noticeable differences between the sequencing platforms. As expected, the quality score profiles along the reads differ, reflecting differences in the base calling and quality score estimation between the instruments. However, GATK’s Base Quality Score Recalibration was effective in minimizing these differences (Supplemental Figure S2). Still, the drop in quality score at the end of Read2 on HiSeqX was more pronounced than on the NovaSeq. We noticed slightly larger number of long homopolymers on the NovaSeq instruments (Supplemental Figure S3). We computed for each read the longest stretch of each possible base and summarized the results in Figure 2. In both Read 1 and Read 2, NovaSeq instruments produced more stretches of G’s than HiSeqX, which we attributed to an artifact resulting from the fact that G is detected as the absence of signal in the 2-color chemistry of the NovaSeq platform. Although less pronounced, we also detected this effect for stretches of A’s (detected by the joint signal of both colors), especially in Read 2 and the inverse effect for C and T bases (each detected by the red and green signal respectively). We believe that some of the patterns, illustrated in Figure 2 (such as the bump of stretches of 81 G’s in Read 1) are also due to differences in the base calling algorithms.

**Figure 2:**
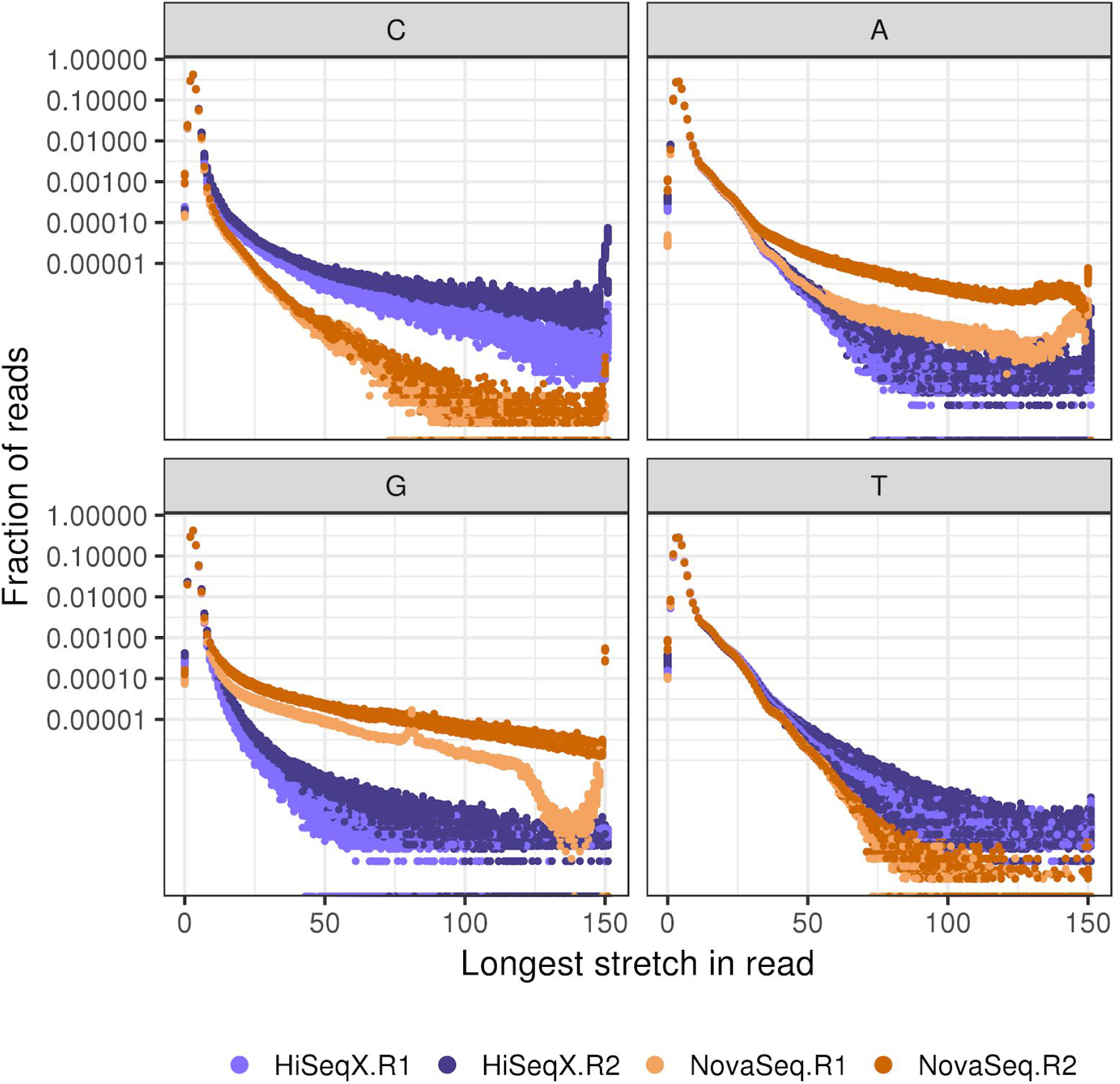
Distribution of length of longest stretches of a nucleotide in HiSeqX and NovaSeq, Read 1 and Read 2 FASTQ files. Each dot represents fraction of reads in a single FASTQ file. Fraction of the total number of reads is represented in log-scale.

### Alignment-level comparison

The mean coverage of the tumor cell lines ranged from 80X to 278X for the tumor cell lines, and from 42X to 180X for the normal cell lines. The alignment rate was very comparable between the two platforms and always superior to 99.5%. The percentage of reads marked as duplicates was higher on HiSeqX (mean 11.25%) than NovaSeq (mean 6.6%), despite the deeper coverage of the NovaSeq samples. This was unexpected, because we usually see slightly higher duplication rates on NovaSeq as compared to HiSeqX. We attributed this observation to differences in loading concentration between the HiSeqX and the NovaSeq flow cells as this generally correlates strongly with the observed duplication rate for PCR-free libraries.

We observed differences between the sequencing platform in the single nucleotide mismatch profiles, where samples sequenced on HiSeqX contained more C>A and T>A mismatches compared to samples sequenced on NovaSeq which had more A>G and T>C (Supplemental Figure S6). HiSeqX data had an average mismatch rate of 0.75% compared to 0.6% in NovaSeq data. Filtering out low mapping quality reads and low quality bases reduced most of the sequencing platform based differences, leaving higher T>G and lower C>T and G>A mismatches in NovaSeq samples (Figure 3A). The overall mismatch rates in the two platforms after quality filtering were very similar, 0.24% in NovaSeq and 0.23% in HiSeqX.

**Figure 3:**
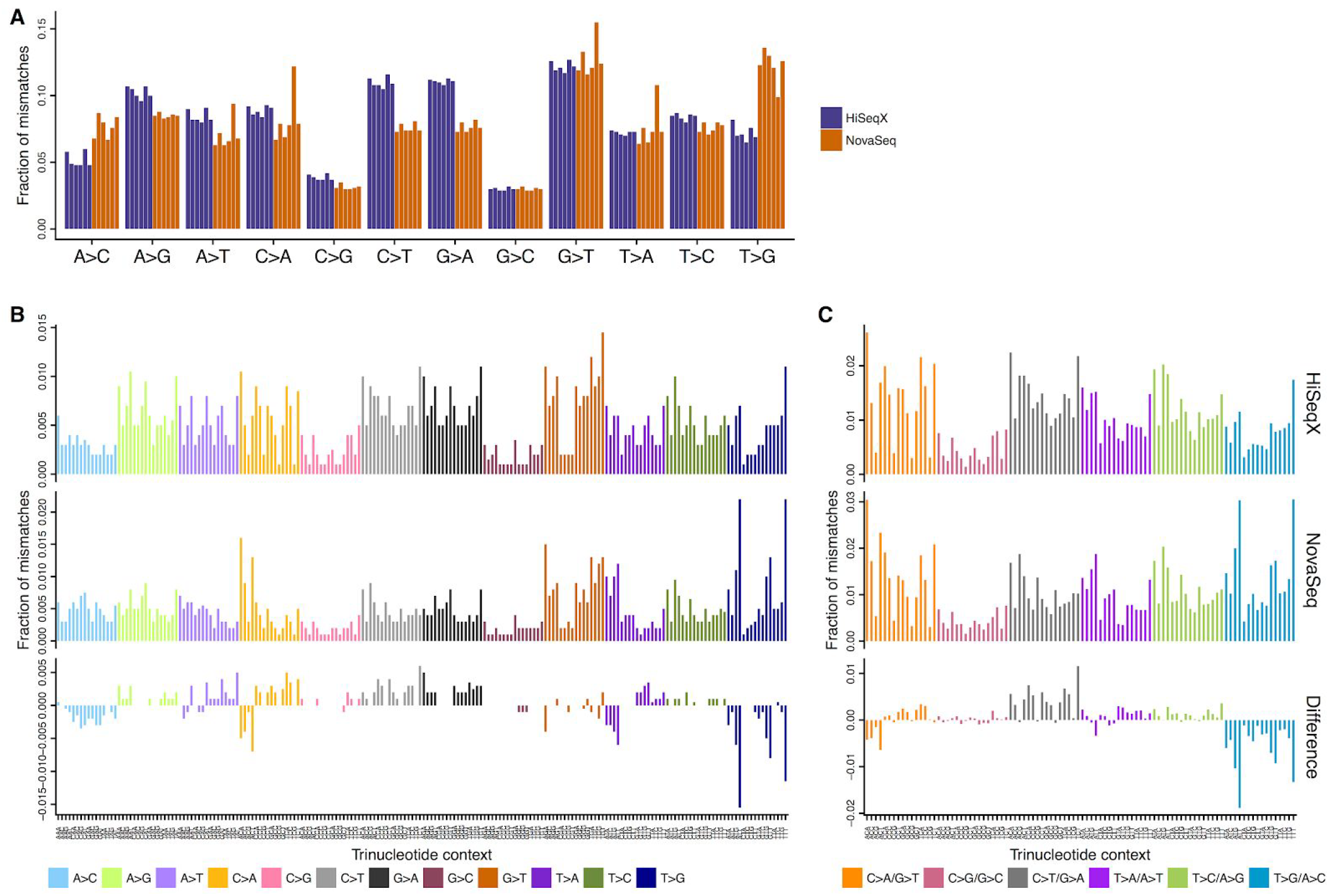
(A) Single nucleotide mismatches by type in samples sequenced on NovaSeq and HiSeqX, with MQ≥10 and BQ≥10 cut-offs. Each bar represents a single sample and is colored based on sequencing platform. (B) Average mismatch rates for bases with MQ≥10 and BQ≥10 across the 6 cell line samples for each mismatch type per trinucleotide for HiSeqX (top row), NovaSeq (middle row) and difference between HiSeqX and NovaSeq (bottom row). (C) Same as (B), but with mismatch types categories collapsed with their respective reverse complements.

The mismatches were further split based on their trinucleotide context. Figure 3B shows the fraction of mismatches averaged across all samples. We noticed that T>G mismatches in NovaSeq samples were predominantly found in the trinucleotide contexts of A[T>G]T, G[T>G]T and T[T>G]T. In order to compare with the somatic mutational profile, all mismatches were collapsed to the 6 mismatch types (C>A, C>G, C>T, T>A, T>C, T>G) (Figure 3C). Even after collapsing the mismatch types, we saw more T>G/A>C mismatches in the NovaSeq data. The difference in the mismatches between HiSeqX and NovaSeq, for all samples is shown in Supplemental Figure S7, and for the collapsed mismatch types in Supplemental Figure S8.

### Variant-level comparison

In order to compare the variants called in HiSeqX and NovaSeq data for the same tumor-normal pairs, we first downsampled all the tumor samples to 80X coverage and the normal samples to 40X coverage and ran our variant calling pipeline. For all PASS-filtered somatic variants called by the pipeline, the concordance between the platforms ranged from 81 to 92% for SNVs, 45 to 58% for indels, and 50 to 77% for SVs (Figure 4). When comparing the high confidence variants, the concordance was much higher: 90 to 94% for SNVs, 87 to 94% for indels and 81 to 88% for SVs. This clearly illustrates the advantages of using multiple callers and evidence from orthogonal strategies to reduce false positive calls, particularly for indel calling. We also compared two NovaSeq replicates of COLO-829 and COLO-829BL sequenced at 80X and 40X respectively, on distinct lanes of the same sequencing run, and found that the intra-run variability is comparable to the inter-platform variability (93% for SNVs, 91% for indels and 83% for SVs).

**Figure 4:**
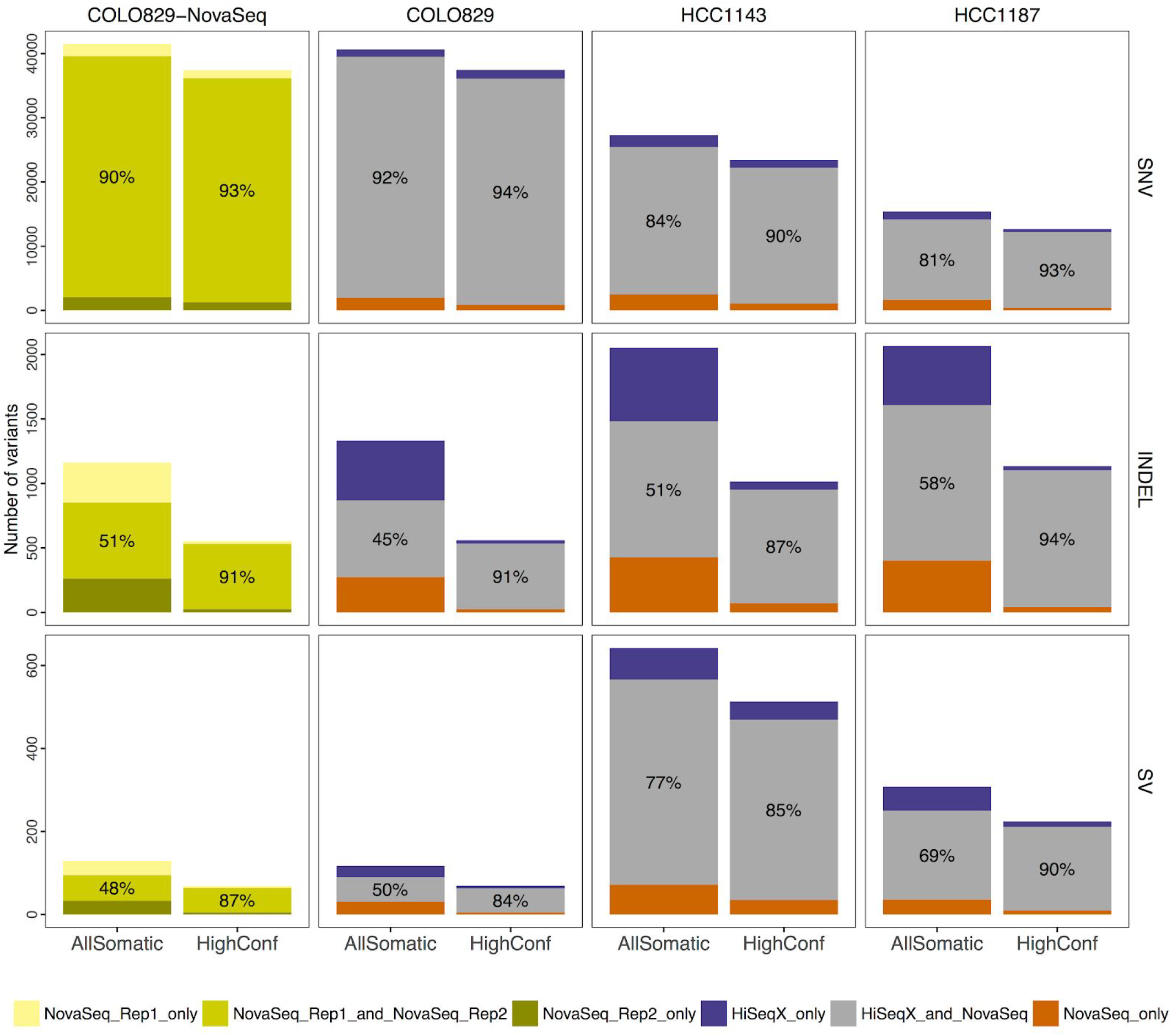
Intra-run and inter-platform concordance of somatic variants. The first row corresponds to SNVs, second row to Indels and third row to structural variants. The first column corresponds to comparisons between two replicates of COLO-829 NovaSeq data (created using reads from mutually exclusive lanes) and indicate the within-platform intra-run variability. Columns 2-4 show comparisons between platforms for the three cell lines. Orange bars (resp. purple) represent the number of variants called uniquely in the NovaSeq runs (resp. HiSeqX) and the grey bars correspond to the variants called in both samples. We indicate the results for all PASS variants from our pipeline (AllSomatic) and for the variants we classified as high confidence (HighConf). All tumors were downsampled to 80X and normals to 40X mean coverage for these comparisons.

Focusing on the discordant calls, we observed that most of the somatic SNVs identified by one platform but not the other are observed at low allele frequency (<10%), indicating that they might have been missed by insufficient coverage and sampling differences, or be false positives introduced by sequencing artifacts (Figure 5B and Figure 5D). We also observed that the mutation spectrum for discordant calls was very different from the concordant calls (Supplemental Figure S5, Figure 5), with relatively large number of T>G mutations among the variants unique to the NovaSeq instrument (Figure 5). This is in agreement with the higher T>G mismatches, especially in A[T>G]T, G[T>G]T and T[T>G]T context, seen in NovaSeq data (Figure 3; Supplementary Figure S5). We did not see the same trend when comparing high confidence variants (Supplemental Figure S10). We therefore think that a lot of these T>G variants called only in NovaSeq data may in fact be artifactual calls that could be filtered out if we include more NovaSeq samples in our panel of normals. Our PON predominantly consisted of HiSeqX samples and may therefore be better at removing HiSeqX-specific artifacts. For example, we saw that a lot of T[C>A]A SNV calls unique to HiSeqX (Supplemental Figure S10) were filtered because of the panel of normal filtering step, and therefore were not seen in our final callsets.

**Figure 5.**
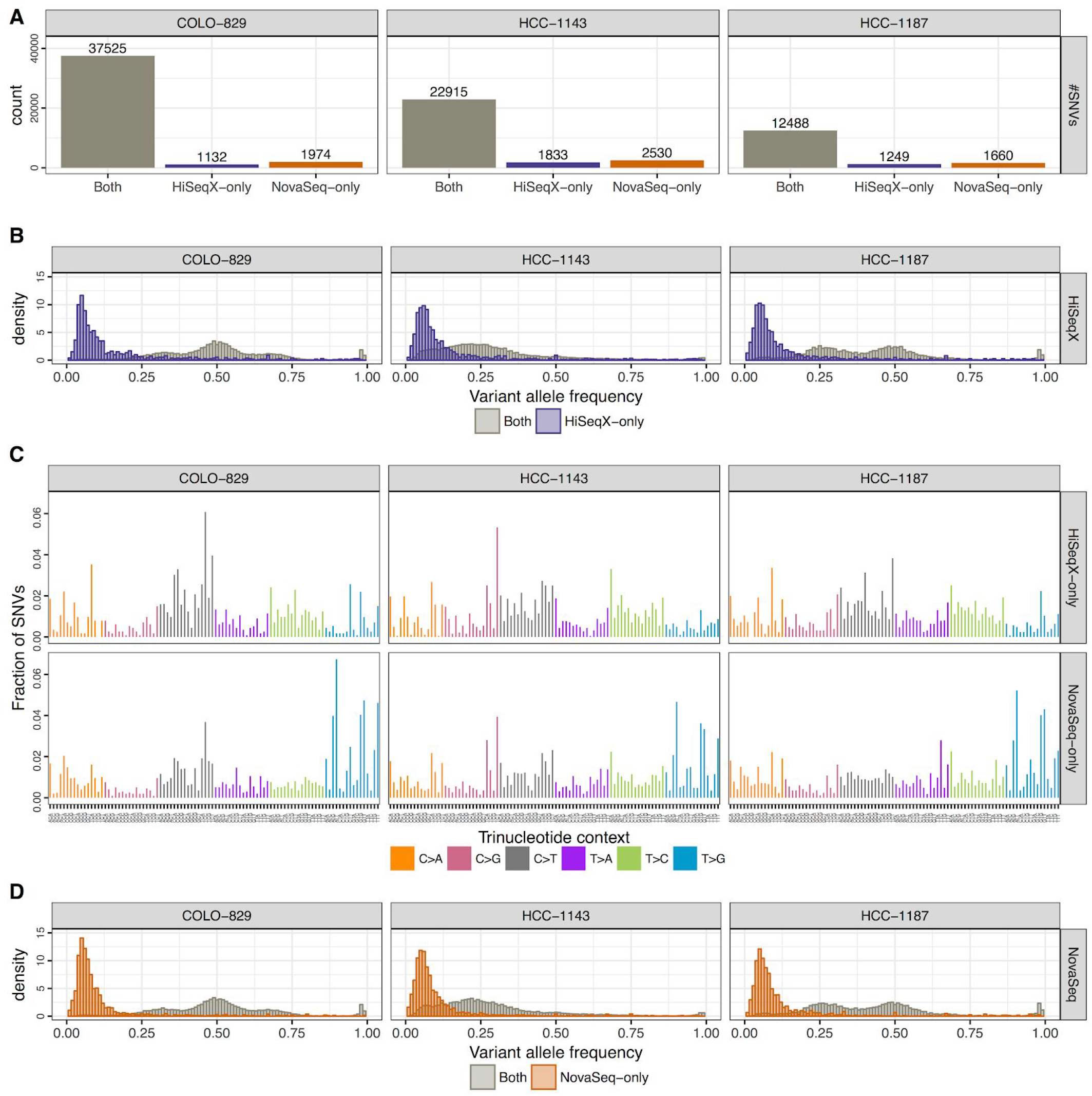
Allele frequency and mutational spectrum of discordant SNVs between HiSeqX and NovaSeq. Panel A shows the number of SNVs that were called in both NovaSeq and HiSeqX data, only in HiSeqX data and only in NovaSeq data. Panel B shows the allele frequency of the variants called only by HiSeqX in purple, and for reference the allele frequency of variants called by both platforms. Panel C shows the decomposition in trinucleotide contexts of the variants called uniquely by each platform. Panel D is similar to Panel B but for variants uniquely called by NovaSeq.

**Figure 6:**
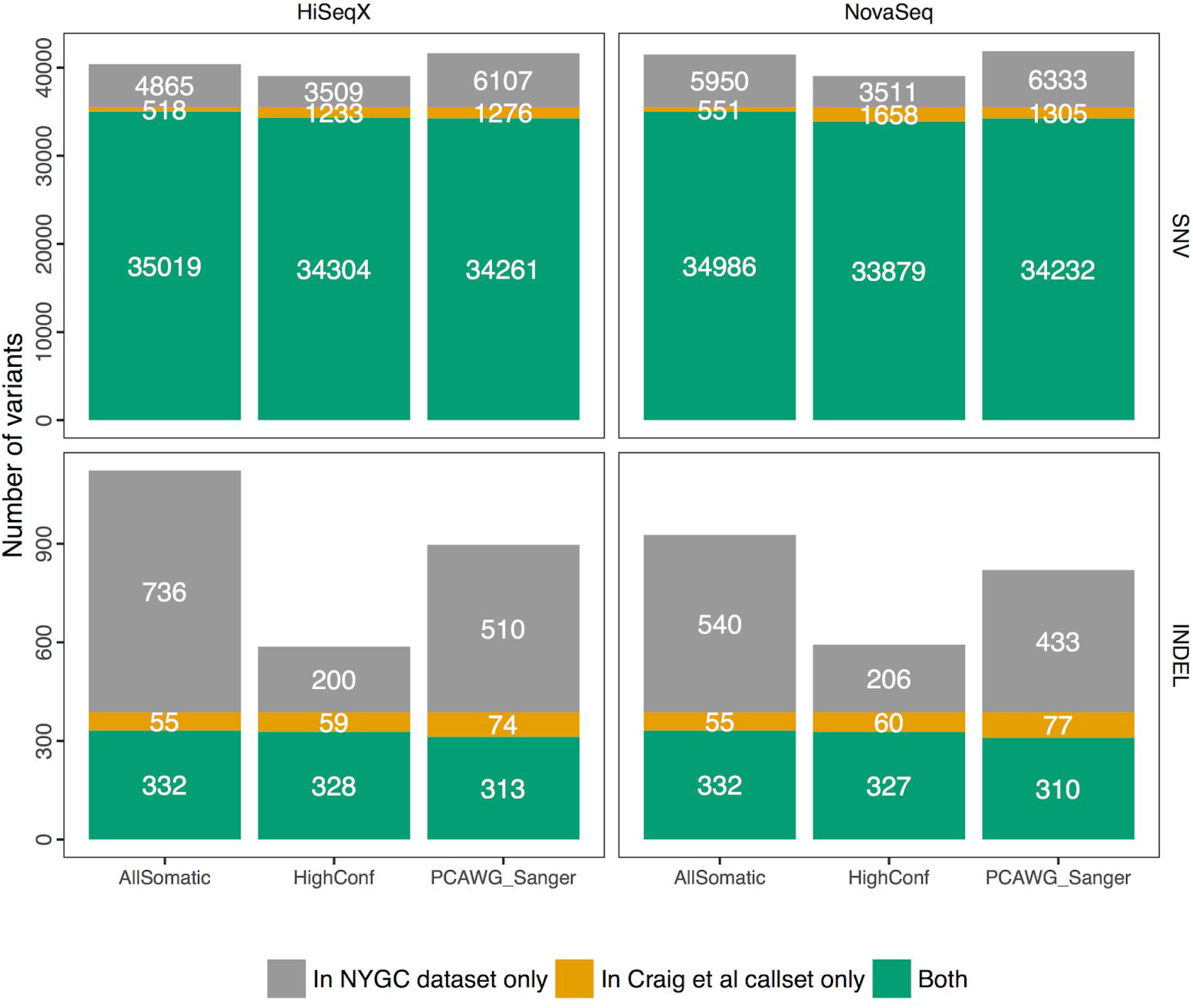
Comparison of somatic variants called on HiSeqX and NovaSeq COLO-829 tumor/normal data downsampled to 80X/40X to the Craig et al. reference dataset.

### Comparison to a reference callset and an alternative somatic pipeline

The three cell lines we sequenced have already been extensively studied and sequenced by other groups. In particular Craig et al. sequenced different passages of COLO-829 in three different centers (TGen, BCSGSC and Illumina) and established a somatic reference dataset for SNV and indels from the consensus of their pipelines. This dataset also provided copy number gain/loss information for 6,586 genes. In Pan-Cancer Analysis of Whole Genomes (PCAWG) project, three best-practice pipelines were developed from the participating institutions and made accessible in Docker containers [33]. Of the three pipelines, we ran the pipeline from Sanger institute on COLO-829 samples sequenced on HiSeqX and NovaSeq at NYGC. The other two pipelines from DKFZ and Broad institute were not readily implementable, due to assumed dependencies in those workflows. The PASS variants from the PCAWG Sanger pipeline were compared against the Craig et. al reference dataset.

Overall, we saw that our pipeline called fewer SNVs than Sanger pipeline, and yet we called over 98% of the Craig et al. SNVs compared to around 96.4% called by the Sanger pipeline. Our pipeline called more indels compared to the Sanger pipeline, but we had far fewer indel calls in the high confidence callset. We called around 86% and 84% of the Craig et al. indels in our All Somatic and High Confidence callsets, compared to 80% called by the Sanger pipeline, suggesting that our pipeline may be more sensitive.

We explored the sources of discrepancies between our callset and the reference dataset established in Craig et al. and represented the different categories in Supplemental Figure S11. Almost half of the variants absent from our final callset were in fact called as PASS-filtered by at least one of the callers, but were removed from our AllSomatic list due to PON filtering. Some of the discordant variants were in the raw callsets of one or more variant callers, but did not pass the caller-specific filtering. Other sources of discrepancies were evidence of the variant allele in the normal sample, low coverage or low variant allele frequency in the tumor sample. While there were some differences between SNV and indel calls between the two pipelines, we found that the CNV recall was very similar between the two pipelines based on a gene-level comparison (99.8% recall for both our pipeline and the Sanger pipeline).

Overall, despite some discrepancies, we are confident about our callset and believe that the PON filtering is a powerful method to remove technical artifacts. It is also entirely possible that some somatic variants were different between the cells used in the reference dataset and the ones used in this work.

### Recall and Precision at Various Purities

One frequent concern in cancer genomics is that tumor samples are always heterogeneous, composed of tumor cells, stromal contamination and normal cells. Since one goal of a somatic pipeline is to establish the catalog of the somatic mutations occuring in the tumor cells, it is important to take into consideration the composition of the sample, usually summarized as the “tumor purity”. Using the two most deeply sequenced cell lines, we simulated lower purity tumor data and evaluated the performance of our SNV/indel pipeline at different purities by comparing to the results obtained with high coverage (see Methods). We observed that precision remained good for high-confidence SNVs and indels even at low purity, but that recall decreased rapidly with purity lower than 50% for the highly rearranged HCC-1143, and below 25% for COLO-829, which had comparatively fewer chromosomal abnormalities (Figure 7A, Supplemental Figure S13). We conducted a similar type of precision/recall analysis for CNVs (Figure 7B), comparing calls across samples at the base-pair level and found that for amplifications, the recall tended to decrease more gradually as purity decreased but for deletions there was a sharp drop-off below 50% purity. This is mainly because deletions occupy few copy number states (e.g. 0 or 1 copy for a diploid genome), whereas amplifications can have higher copy number states that will still be captured at lower purity, as is the case with HCC-1143. Precision remained relatively high at different purities for both amplifications and deletions; however, for deletions it dropped off at around 25%, as very few calls were being made at this level of purity.

**Figure 7.**
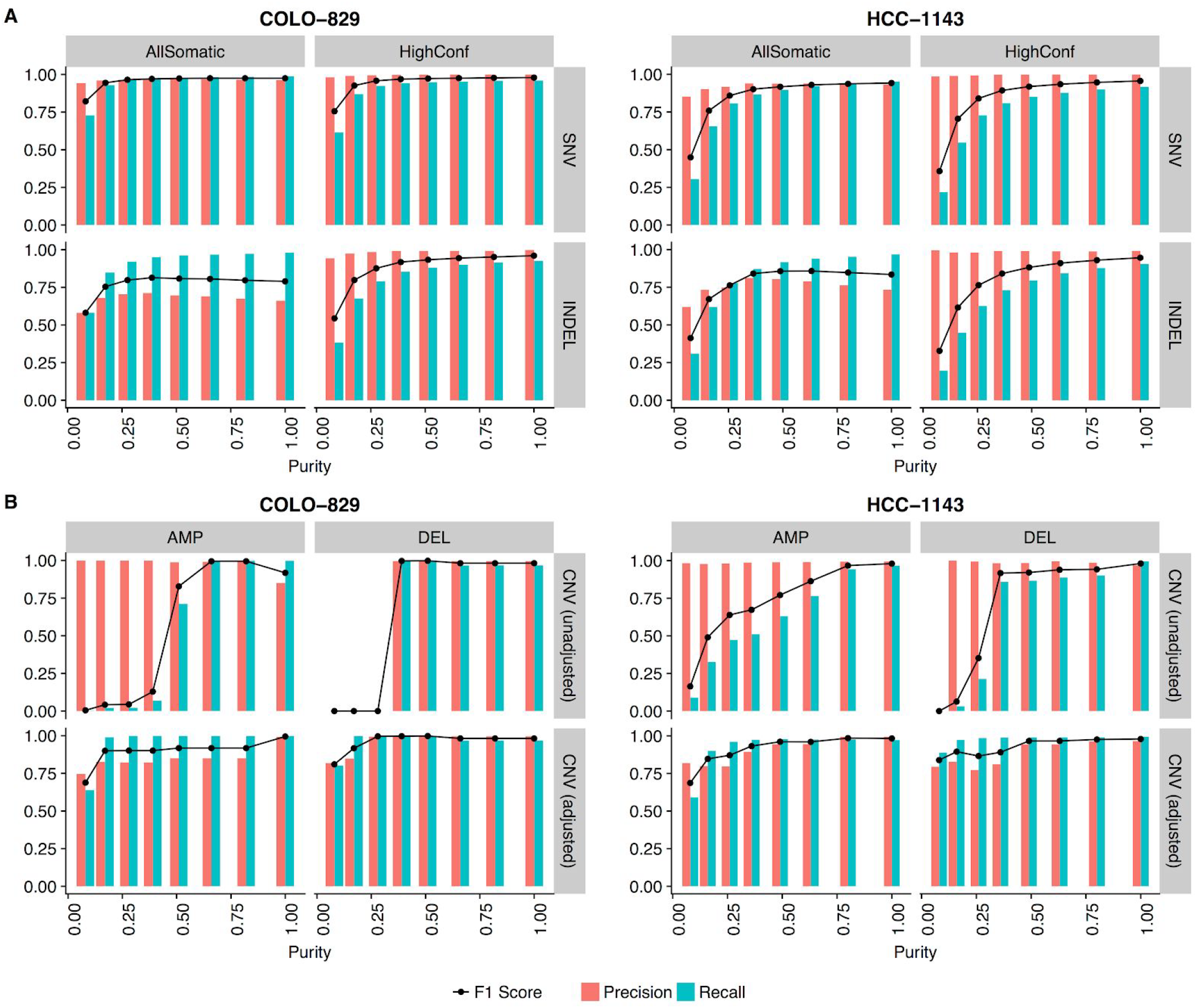
Precision, recall and F1 scores at different simulated purities for (A) SNVs (top) and Indels (bottom), and (B) CNVs without (top) and with (bottom) adjustments of log2 values for purity and ploidy.

For low purity samples, some of the false negative CNV calls could be attributed to the chosen thresholds for categorizing amplifications and deletions, and the fact that BIC-seq2 outputs log2 ratios that are not adjusted for purity and ploidy. We wanted to investigate how purity/ploidy adjustment could rescue true events missed in low purity samples. For this, we based our purity estimates off of the fraction of reads we mixed from the tumor and normal sample to create these low purity samples, which should represent the true purity (see Methods). Using these estimates, the CNV log2 values were adjusted. We found that the recall was much higher than the original unadjusted data, but precision gradually decreased as the purity dropped. At the lowest purity level, precision and recall dropped, which was likely a result of the different segmentation at this purity level. The ability to capture CNV calls in lower purity samples by adjusting the log2 values based on purity/ploidy is very useful, but requires correct estimation of purity and ploidy for the tumor sample.

Therefore, we evaluated CELLULOID [32] and HATCHet [34] for their ability to correctly estimate the purity and ploidy values for our purity ladder samples (Figure 8). Both tools use read depth information at germline heterozygous sites to infer the tumor purity/ploidy. In particular, HATCHet can be run in multisample mode, which can leverage information from high purity samples to infer the purity/ploidy of low purity samples from the same individual. We ran HATCHet in both single sample and multisample mode. We found that both tools tend to perform well above 50% purity. Furthermore, HATCHet (in multisample mode) and CELLULOID can both give close estimates of the purity/ploidy for much lower purity samples. CELLULOID estimates only seemed to drop-off for samples with a purity lower than 12.5%, while HATCHet in multisample mode produced a good estimate at this low purity level.

**Figure 8:**
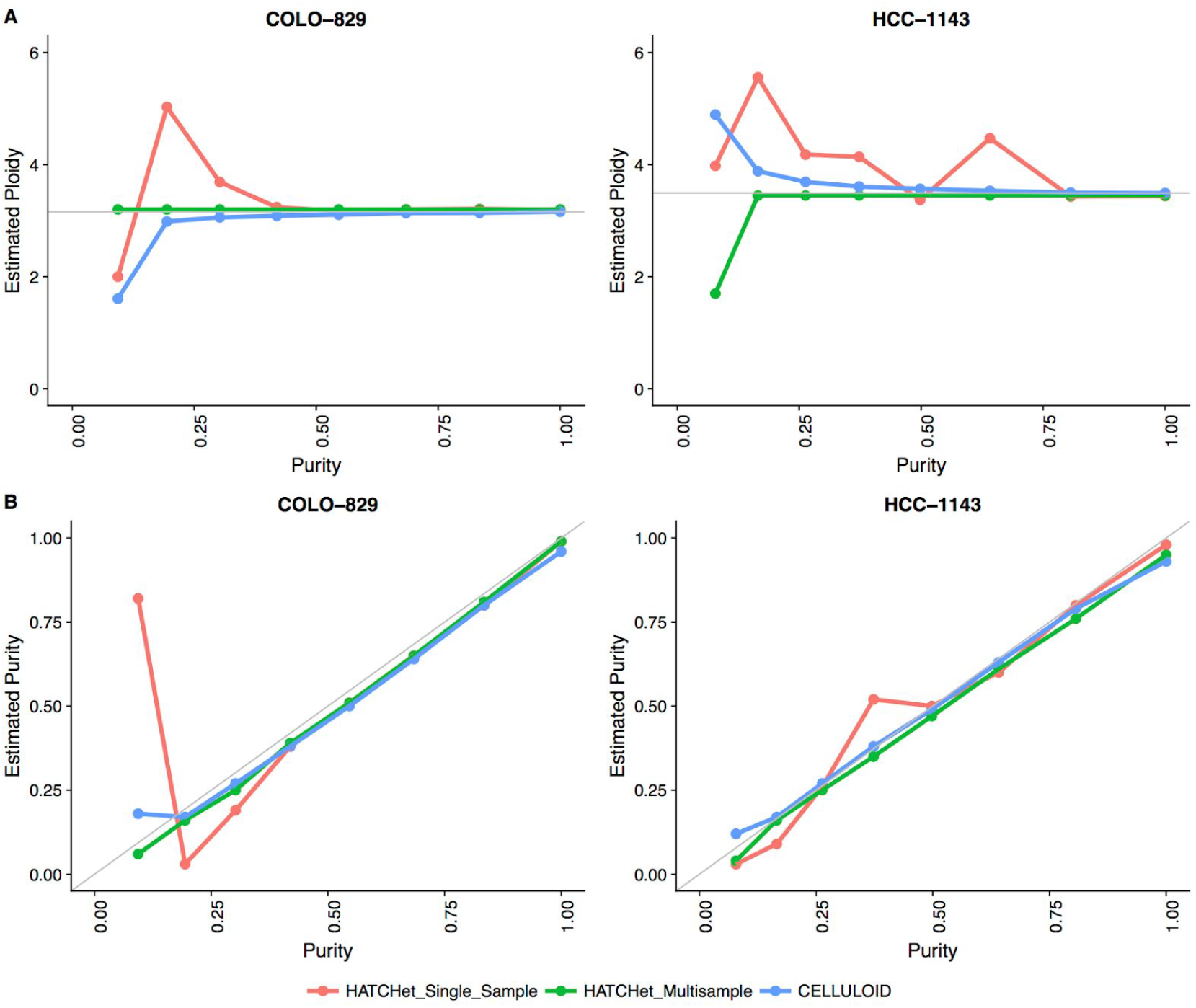
Ploidy and Purity Estimation of Cell Line Purity Ladder using CELLULOID and HATCHet in single-sample and multi-sample mode

Further, we looked into the log2 values of the events that were captured at lower purities and their adjusted log2 values based on CELLULOID and HATCHet single sample estimates of purity and ploidy (Supplemental Figure S12). In the low purity samples, we were able to identify many of the events that were originally lost at this purity level. However, it was also apparent that the CNVs being captured at the lowest purity level did not necessarily resemble the high purity CNVs. This is another example of how differences in segmentation in lower purity samples can have an impact on recapturing CNV calls.

## Discussion

Cancer cell lines are useful models for studying cancer biology. They are widely available, easy to propagate and composed of a relatively homogeneous population of cells, making them extremely valuable for advancement of tools and methods for cancer genomics. However, they are imperfect models and do not represent the entire complexity of real tumor samples. They may also contain unique genomic features needed for immortalization and in vitro growth. Here, we used 3 cancer cell lines for benchmarking purposes and share our high-quality callset to the genomics community. We plan to keep using this data to test novel variant callers and may resequence these cell lines with novel sequencing technologies (such as long read technologies). Here, we demonstrated using these cell lines the existence of systematic differences between the reads produced by HiSeqX and by NovaSeq. The patterns we identified will need to be taken into account to fully exploit the signal produced by the sequencers. For instance, we believe that a deep learning model designed to filter out sequencing artifacts and detect real mutations at very low frequency (such as would be needed for early detection of cancer in liquid biopsy samples) would need to be trained independently for HiSeqX and for NovaSeq, depending on the instrument used for the real-life application of the model. We designed a pipeline for somatic variant calling, composed of multiple softwares for SNV, indel and structural variants. We showed the advantage of using multiple tools to obtain high confidence calls. We showed that with the standard coverage of tumor-normal whole genome sequencing (80X/40X) and our somatic pipeline, the pattern of homopolymer frequency does not translate into systematic biases once multiple somatic callers are applied. We noted a mild enrichment of T>G mutations in the variants called uniquely in NovaSeq and not in HiSeqX data. However, that was not the case when we compared our high confidence variants (those that are supported by multiple callers). Overall, this gives us the confidence to upgrade our sequencing platform to NovaSeq, without any loss of quality (and with a substantial gain in the cost of sequencing and a higher throughput). We demonstrated the importance of filtering recurrent artifacts with a Panel of Normals, ideally composed of a large number of samples from the platforms used to sequence the samples of interest and preferably using the same sequencing protocols. We expect to increase the number of normal samples included in our PON, especially from NovaSeq, as we keep sequencing properly consented samples. We plan to explore refined strategies to filter artifacts based on the allele detected in normal samples rather than, as currently, based on the location in the reference genome. Finally, we used the deeply sequenced libraries to test tools designed to estimate purity and ploidy of tumor samples and showed the importance of incorporating these estimates to improve copy-number detection.

## Conclusion

We present high-quality, deeply sequenced whole-genome data for 3 common cancer lines. We used these samples to study in detail the differences between the two most recent high-throughput sequencers from Illumina, HiSeq X Ten and NovaSeq 6000. We ran these tumor-normal pairs through our somatic pipeline and demonstrated that the inter-platform variability was very similar to the intra-run variability, indicating that the systematic differences between the platforms at the read level are well-handled by the base calling algorithm and by our somatic pipeline. We demonstrated the advantages of combining multiple algorithms to detect SNV, indels and structural variants. We used the samples to study in details the effect of tumor purity on performance and tested tools aiming at estimating purity from WGS data. We show how to use these estimations to recalibrate copy-number events and re-categorize amplifications and deletions.

## Data accessibility

We will deposit the raw genomic data and the somatic variants on dbGAP. The somatic variants for HighCoverage and downsampled 40X/80X are directly accessible on our website^8^. The website also contains the sample reports generated by the pipeline and a link to the Outpost QC interface.

## Acknowledgments

We thank Marcin Imielinski (NYGC-Cornell) for the HCC-1143 and HCC-1143BL cell lines, Simone Zaccaria and Ben Raphael (Princeton University) for their help with HATCHet. We also thank the Computational Biology team at the New York Genome Center for their support and suggestions, Stephen Bavier for his help with the companion website, Kevin McGhee for his help with data repository, Rajeeva Musunuri for the Outpost QC interface, the NYGC Research Computing group for their help with data processing and the NYGC Software Engineering team for the automation of the pipeline.

## Supplemental information

**Supplemental Table 1:** Cell line information from ATCC

| Cell line | Ampule passage number<br>(from ATCC) |
| --- | --- |
| COLO-829 | 10 |
| HCC-1143 | 5 |
| HCC-1187 | 28 |

**Supplemental Table 2:** Alignment metrics and duplication rates

| Sample | Platform | Total Reads | %Aligned Reads | Mean Coverage | %Duplicates |
| --- | --- | --- | --- | --- | --- |
| COLO-829 | HiSeqX | 3,942,506,750 | 99.64 | 166.36 | 11.62 |
| COLO-829 | NovaSeq | 5,111,570,566 | 99.57 | 228.06 | 6.28 |
| COLO-829BL | HiSeqX | 2,125,523,908 | 99.64 | 89.93 | 11.18 |
| COLO-829BL | NovaSeq | 4,062,312,284 | 99.62 | 179.73 | 6.97 |
| HCC-1143 | HiSeqX | 1,928,441,034 | 99.61 | 81.15 | 11.65 |
| HCC-1143 | NovaSeq | 6,310,318,566 | 99.54 | 278.26 | 7.28 |
| HCC-1143BL | HiSeqX | 1,017,638,416 | 99.53 | 42.35 | 11.84 |
| HCC-1143BL | NovaSeq | 3,566,322,944 | 99.63 | 155.73 | 7.47 |
| HCC-1187 | HiSeqX | 1,914,759,882 | 99.69 | 79.80 | 11.35 |
| HCC-1187 | NovaSeq | 2,056,483,546 | 99.71 | 90.35 | 6.43 |
| HCC-1187BL | HiSeqX | 1,016,297,632 | 99.63 | 42.58 | 11.24 |
| HCC-1187BL | NovaSeq | 1,390,489,154 | 99.65 | 61.47 | 6.22 |

**Supplemental Figure S1.**
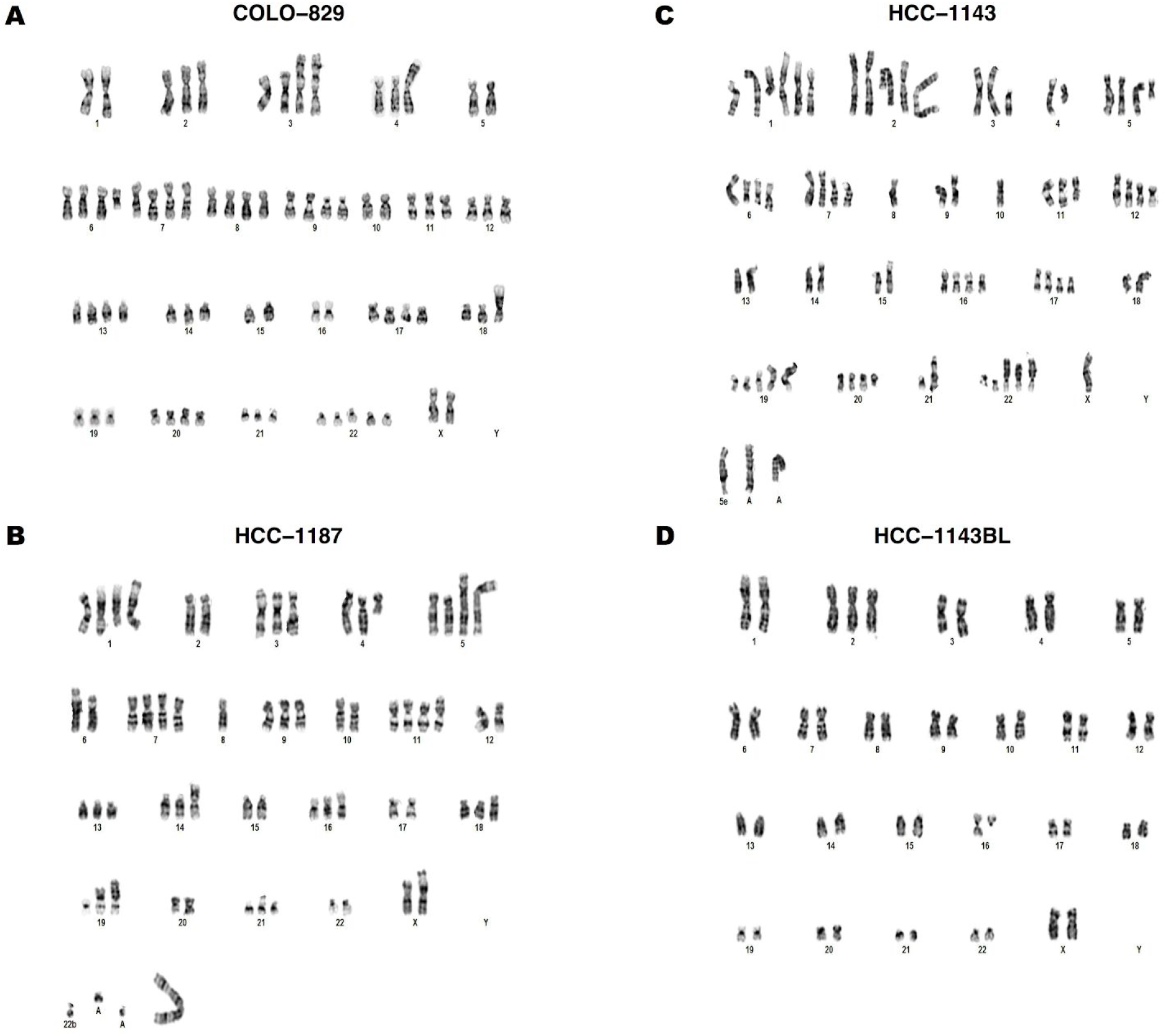
Karyotypes of COLO-829, HCC-1187, HCC-1143 and its associated “normal” cell lines HCC-1143BL. We note some slight differences between the results of the karyotype analyses and the CNV analyses resulting from WGS, possibly due to clonal heterogeneity, technical differences and differences in the level of detection of the technologies. (A) COLO-829: 70~73<3N>,XX,-1,del(1)(q12),+3,der(3)t(1;3)(q12;p25)x2,i(4)(q10),-5,+6,del(6)(q13q25),+7,dup(7)(q32q34) ×2,+8,+9,del(9)(p11.2)x2,-10,+13,-15,-16,+17,der(18)t(1;18)(p21;p11.3),+20,+22,+22,+22 [cp20] (B) HCC-1187: 63~67<3N>,X,add(X)(p22.1),+add(1)(p22),+add(1)(p34),del(1)(q21),del(1)(q32),-2,del(2)(p13p23)x2,+3,del (3)(p13),i(5)(q10)x2,del(5)(q13q33),del(6)(q13),+7,-8,del(8)(q22),-10,+11,add(11)(p15),add(12)(q22),del(13)(q22q32)x3,add(16)(q24),del(17)(p11.2),add(18)(q23),+19,add(19)(p13)x2,-20,add(20)(q13.3),+21,+4~6m ar [cp20] (C) HCC-1143: 74~82<3N>,X,+add(1)(p34),+add(1)(q21),+del(1)(p32p34),+2,add(2)(q31),del(3)(p13),+4,del(4)(q22)x2,+5, del(5)(q13q33),add(7)(q22),del(7)(p13),-8,-10,+11,del(11)(q13q23),del(11)(q23q24),del(12)(q13q22),-14,ad d(14)(p11.2),del(17)(p11.2),+17,add(18)(p11.2),+19,add(19)(p13.3),add(21)(q22),+4~5mar [cp20] (D) HCC-1143BL: 47,XX,+2 [15]/47,XX,+2,del(16)(q12) [5]

**Supplemental Figure S2:**
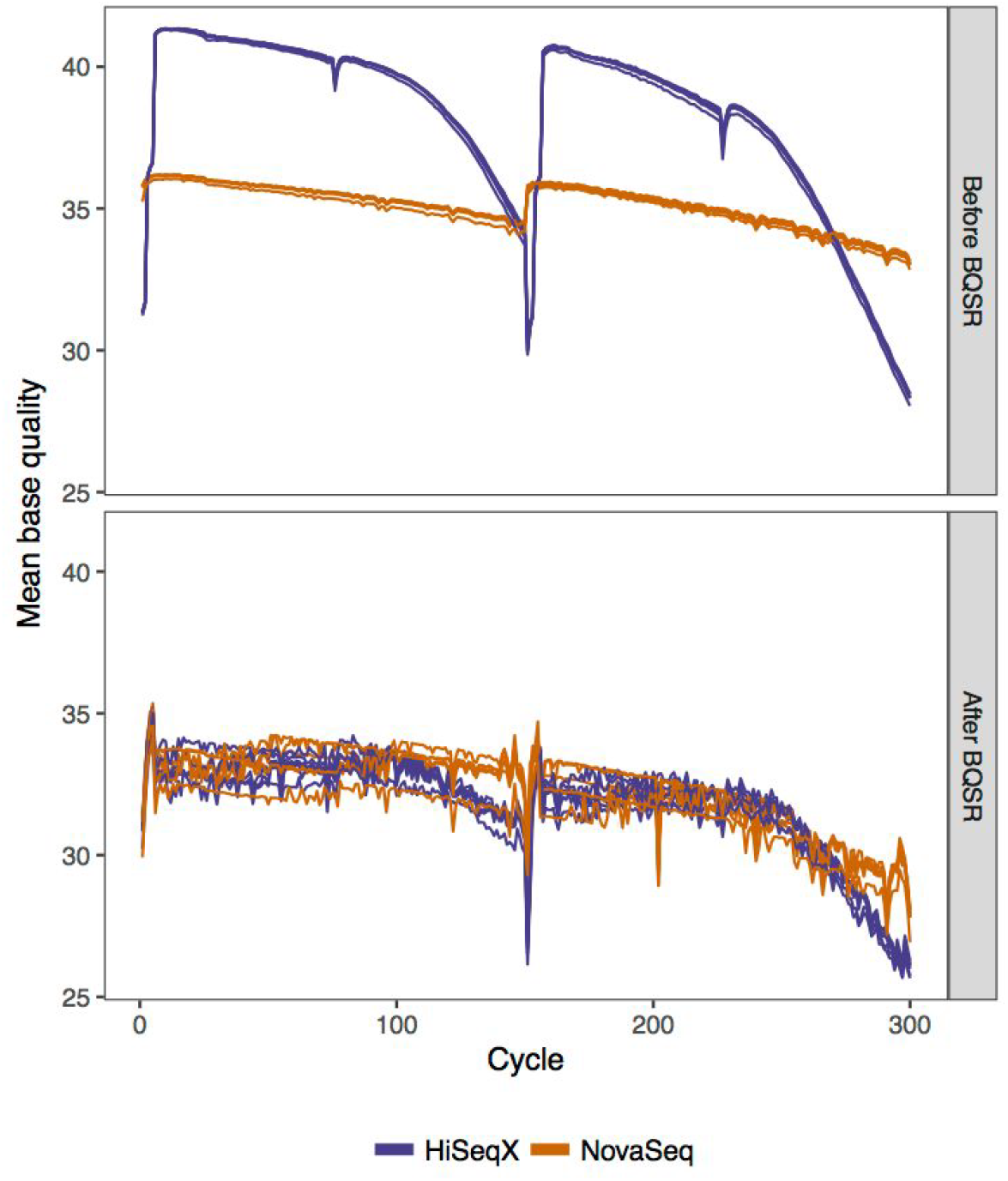
Base quality scores by cycle, before and after BQSR.

**Supplemental Figure S3:**
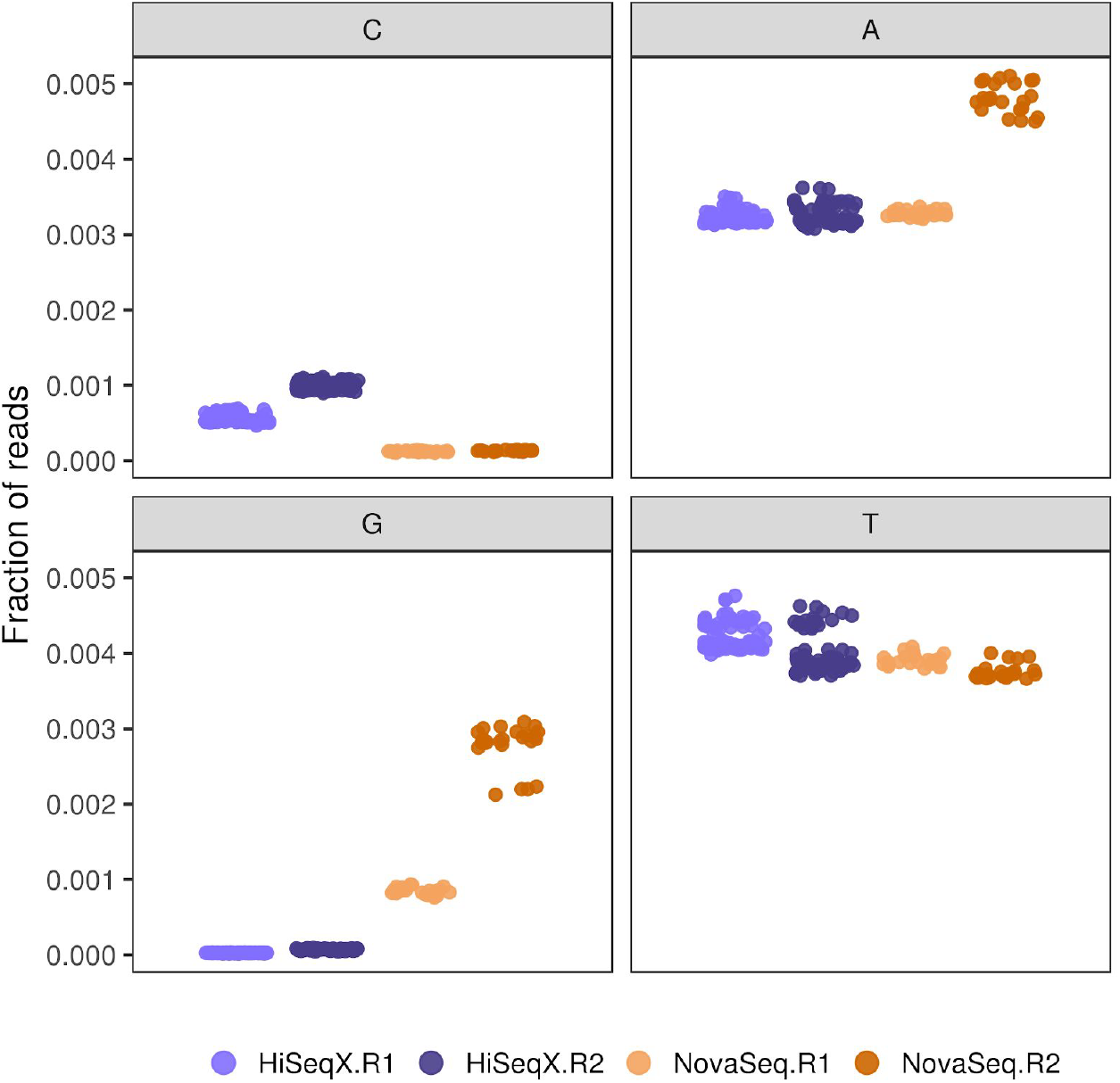
Fraction of total reads containing homopolymer (stretches of 20nt or longer)

**Supplemental Figure S4.**
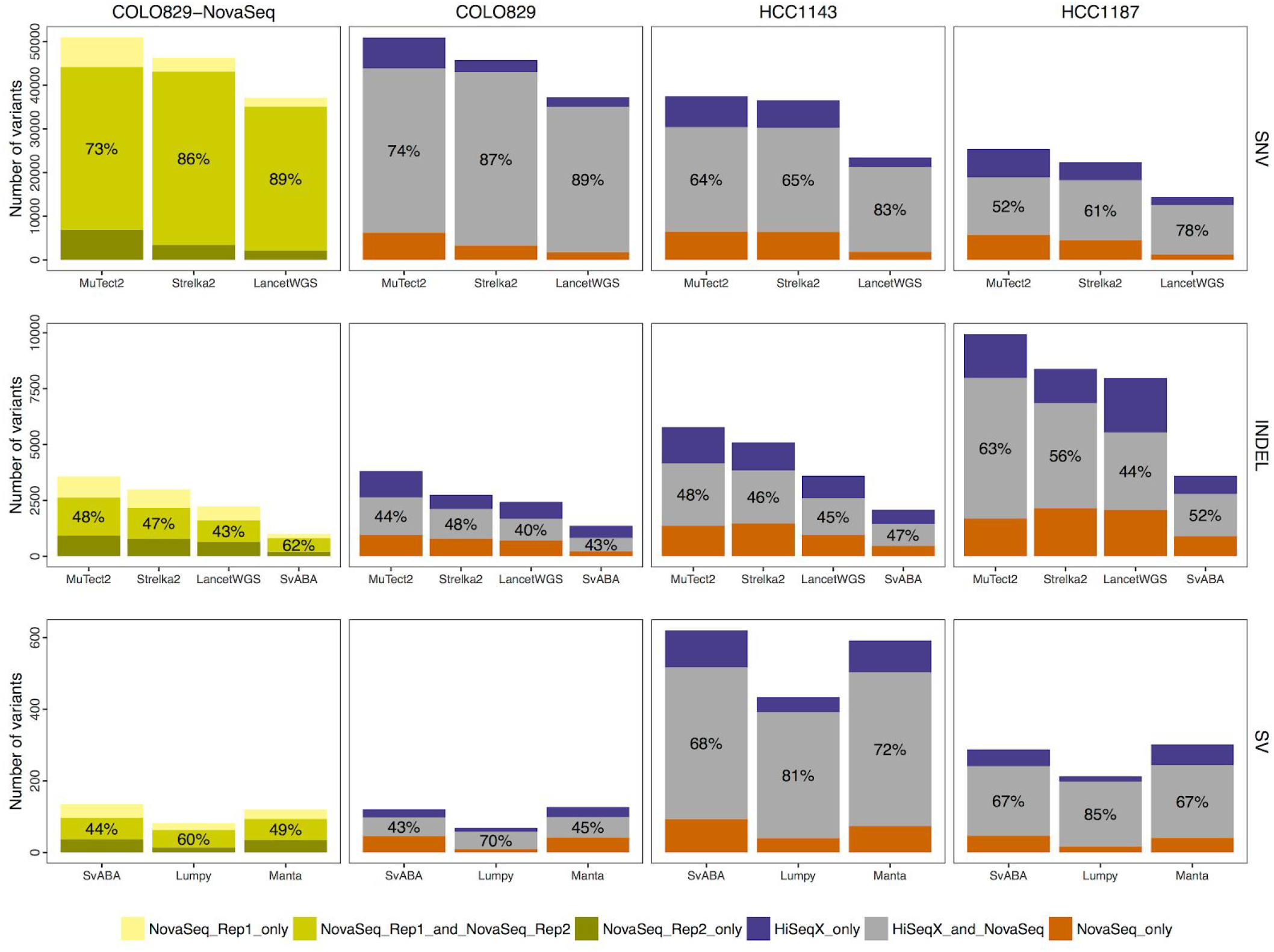
Intra-run and inter-platform concordance of somatic variants called by the different variant callers, similar to Figure 4. Even though Lancet is run in Lancet exonic and validation modes in the pipeline, for this plot, we show the results of Lancet run on the entire genome.

**Supplemental Figure S5.**
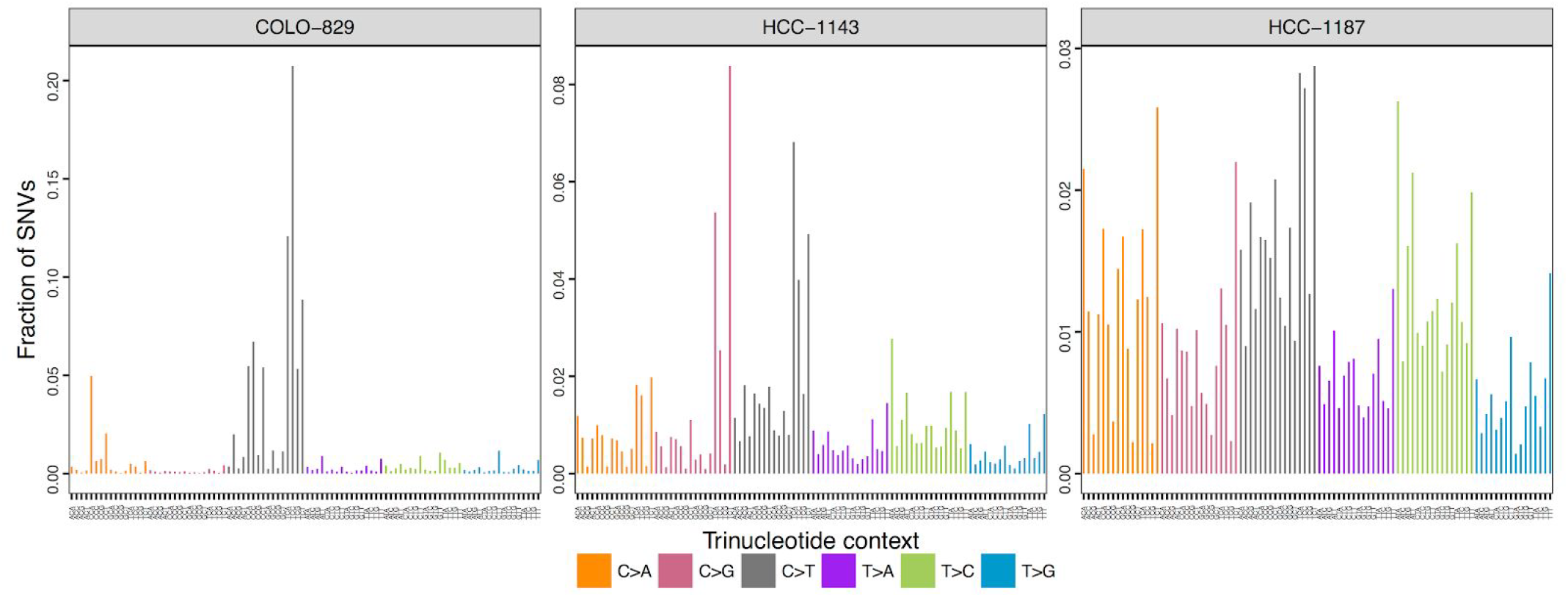
Mutation spectrum of concordant high confidence SNVs between HiSeqX and Novaseq.

**Supplemental Figure S6:**
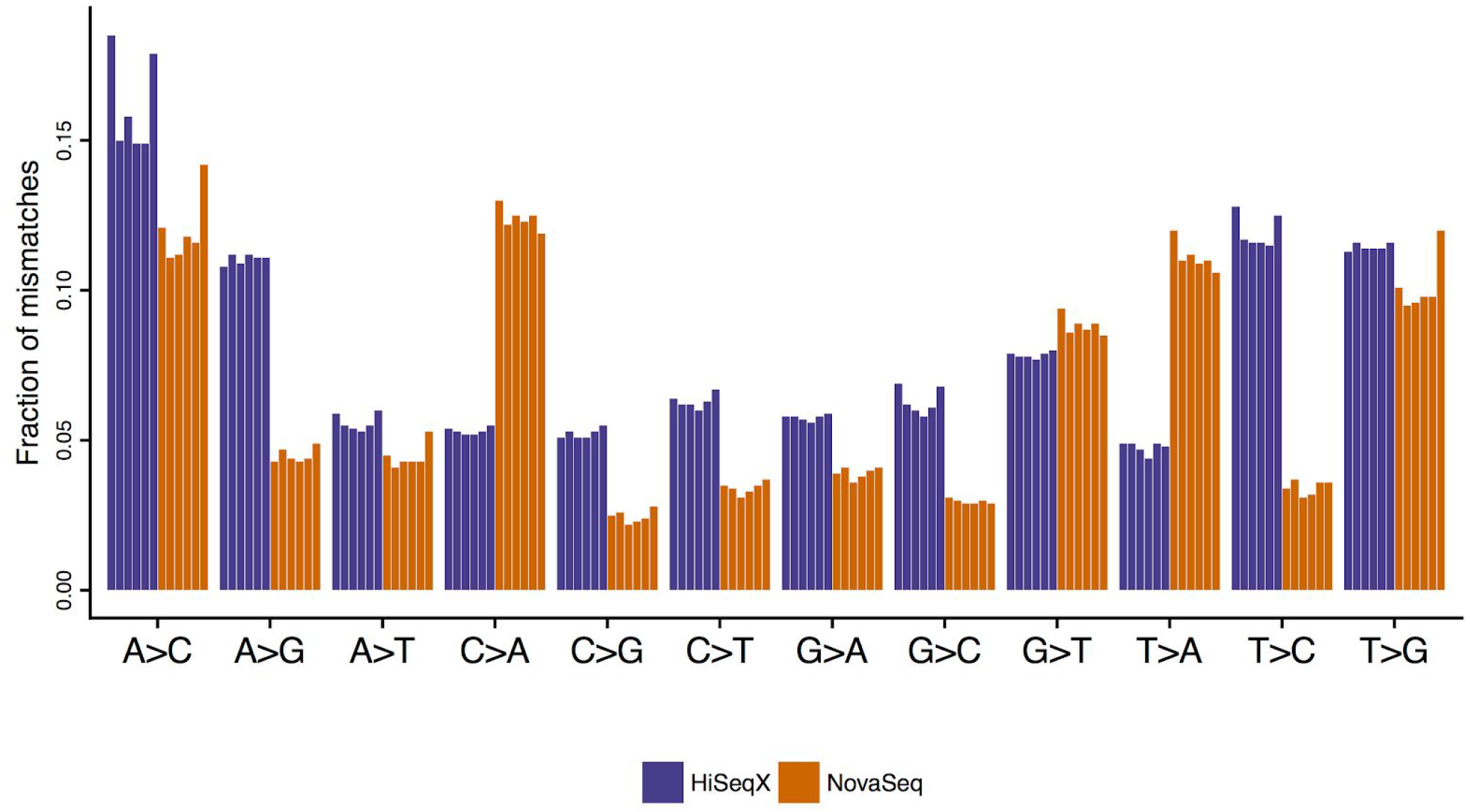
Single nucleotide mismatches by type in samples sequenced on NovaSeq and HiSeqX. We find that NovaSeq had more C>A and T>A mismatches, whereas HiSeqX had more A>G and T>G mismatches. Each bar represents a single sample and colored based on sequencing platform. HiSeqX samples had an average mismatch rate of 0.75%, whereas NovaSeq samples had average mismatch rates of 0.6%.

**Supplemental Figure S7:**
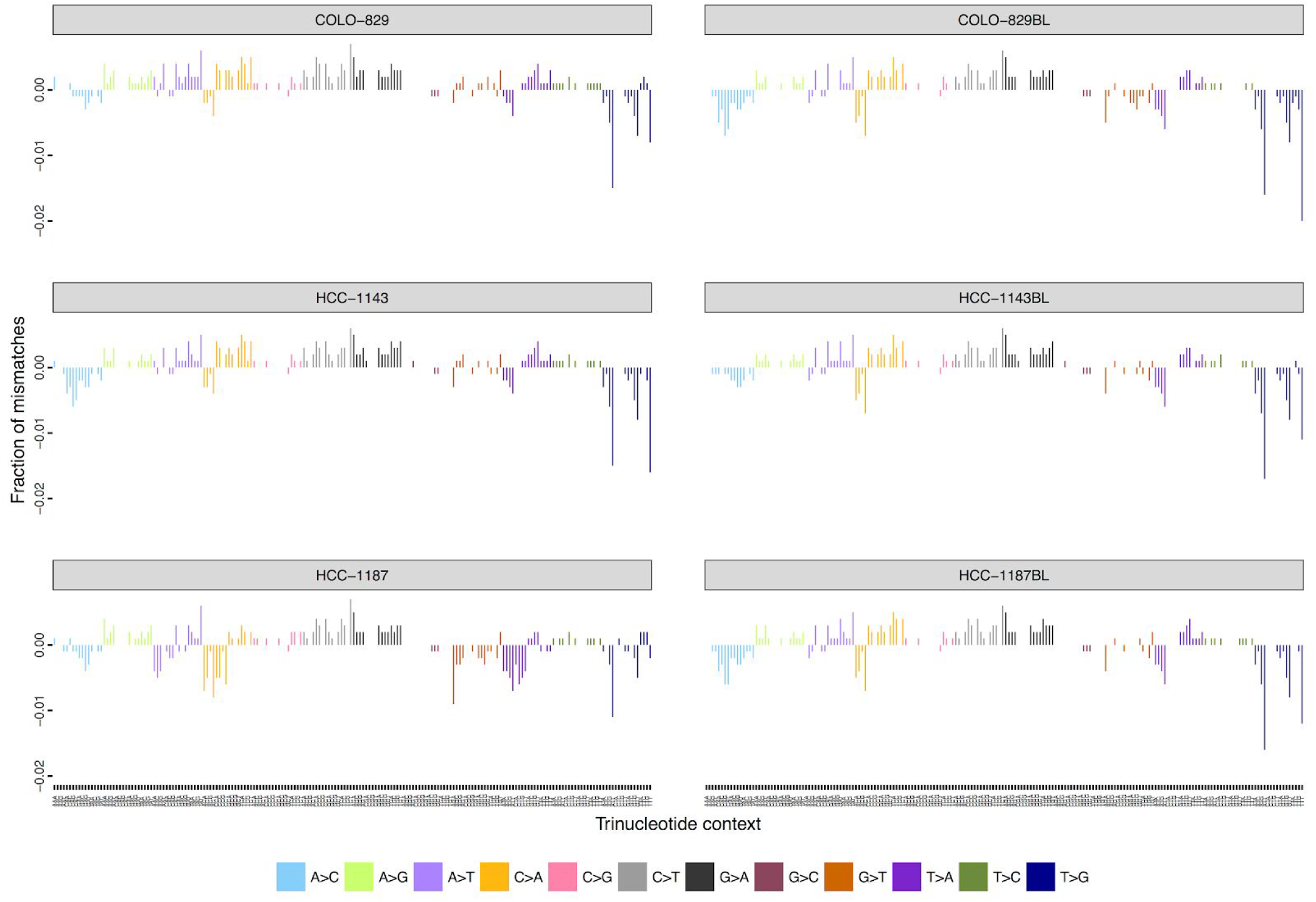
Difference in the fraction of mismatches between HiSeqX and NovaSeq per trinucleotide. Positive values correspond to higher fractions in HiSeqX and negative values correspond to higher fractions in NovaSeq. MQ ≥ 10 and BQ≥10 cut-offs were applied for this calculation. We observed that NovaSeq called more T>G mismatches, especially in A [T>G]T, G [T>G]T and T [T>G]T context.

**Supplemental Figure S8:**
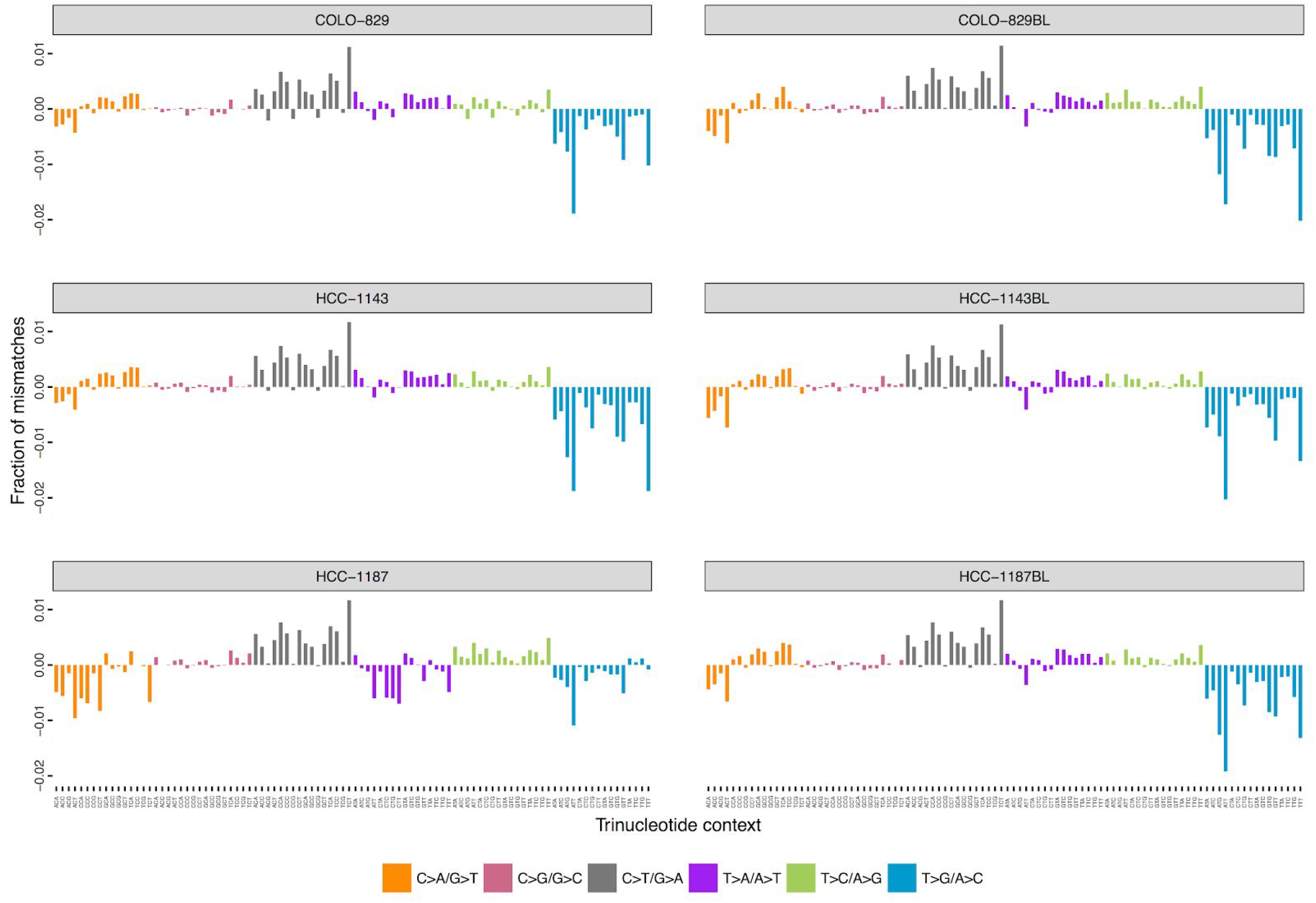
Difference in the mismatches between HiSeqX and NovaSeq per trinucleotide collapsed to the 6 mismatch categories(C>A, C>G, C>T, T>A, T>C, T>G). Positive values correspond to higher fractions in HiSeqX and negative values correspond to higher fractions in NovaSeq. MQ ≥ 10 and BQ≥10 cut-offs were applied for this calculation. We observe that NovaSeq called more T>G mismatches, especially in A [T>G]T, G [T>G]T and T [T>G]T context.

**Supplemental Figure S9:**
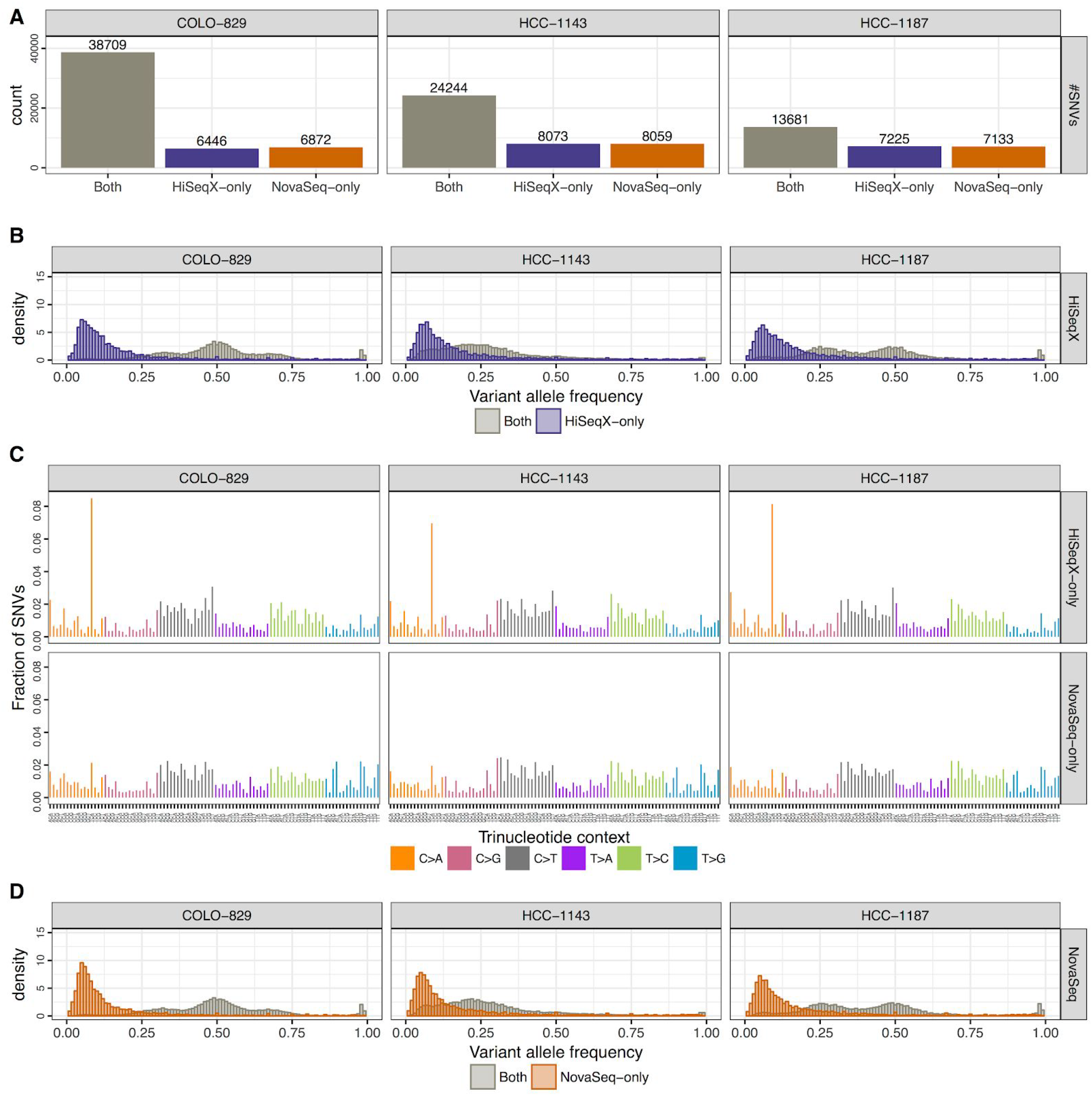
Allele frequency and mutational spectrum of discordant SNVs between HiSeqX and NovaSeq without Panel of Normal filtering. Panel A shows the number of SNVs that were called in both NovaSeq and HiSeqX data, only in HiSeqX data and only in NovaSeq data. Panel B shows the allele frequency of the variants called only by HiSeqX in purple, and for reference the allele frequency of variants called by both platforms. Panel C shows the decomposition in trinucleotide contexts of the variants called uniquely by each platform (top and bottom tracks) and called by both platform (middle track). Panel D is similar to Panel B but for variants uniquely called by NovaSeq.

**Supplemental Figure S10:**
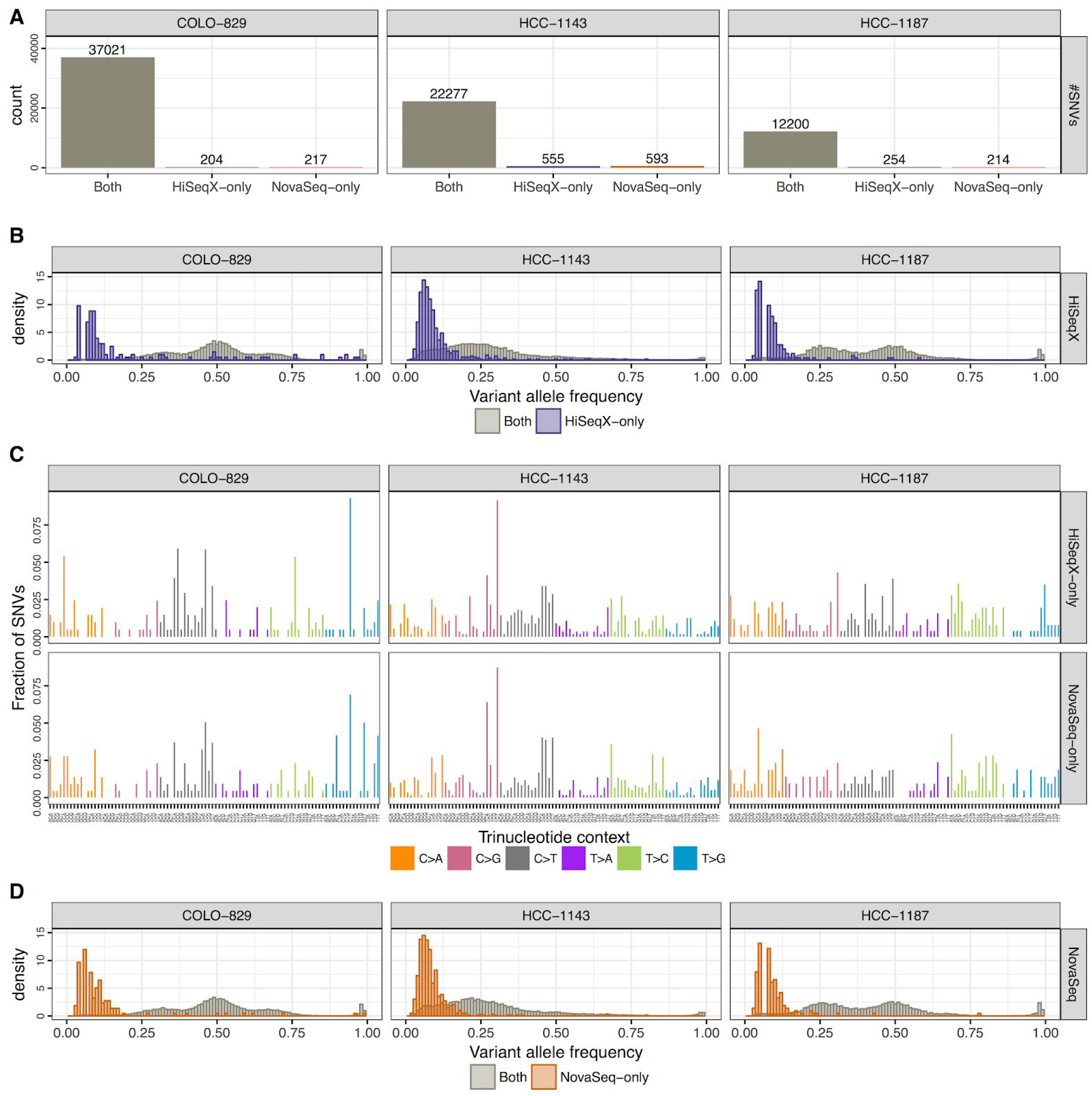
Allele frequency and mutational spectrum of discordant high confidence SNVs between HiSeqX and NovaSeq. Only those SNVs that were in the high confidence callset for at least one of the technologies were used for this. Panel A shows the number of SNVs that were called in both NovaSeq and HiSeqX data, only in HiSeqX data and only in NovaSeq data. Panel B shows the allele frequency of the variants called only by HiSeqX in purple, and for reference the allele frequency of variants called by both platforms. Panel C shows the decomposition in trinucleotide contexts of the variants called uniquely by each platform. Panel D is similar to Panel B but for variants uniquely called by NovaSeq.

**Supplemental Figure S11:**
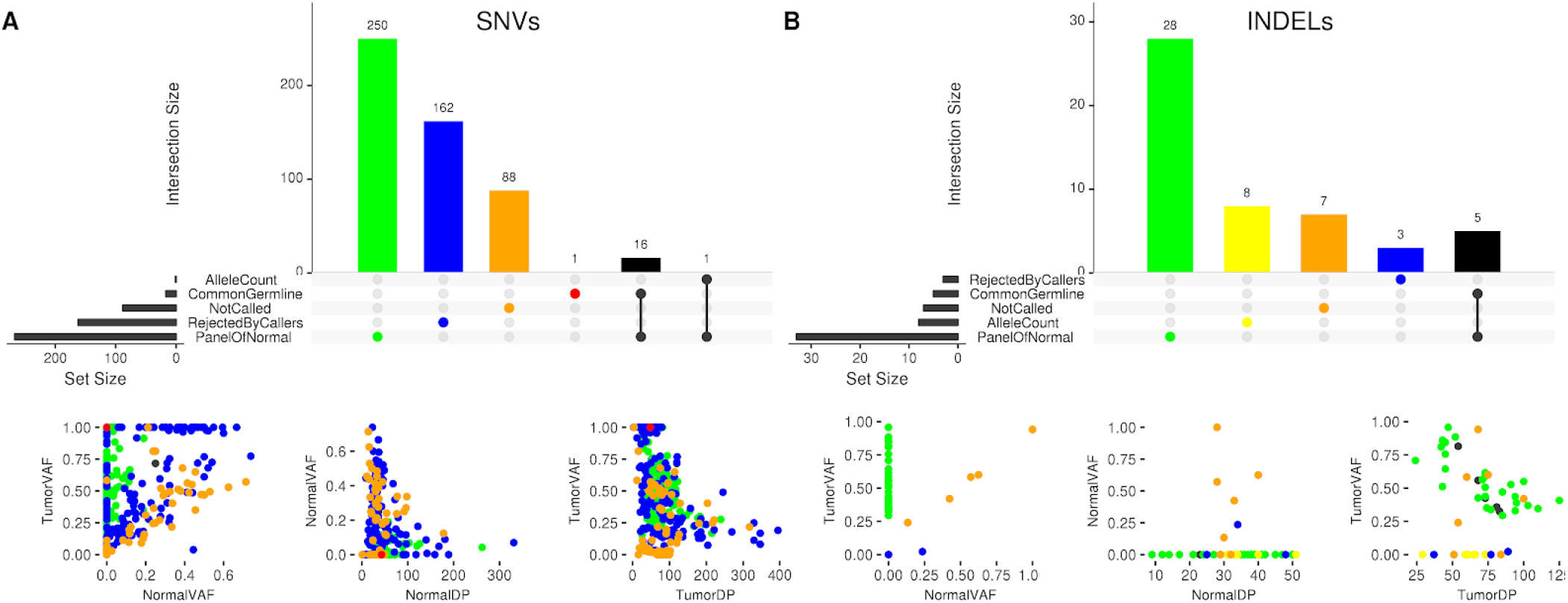
Sources of discrepancies between NYGC callset and the reference dataset established in Craig et al. The figure shows (A) SNVs and (B) Indels from Craig et al dataset that were not called in our AllSomatic callset on the HiSeqX data, and the reasons for rejection or no call: not called by any caller (NotCalled), found only in rejected calls of callers (RejectedByCallers), rejected in Panel of Normals filtering step (PanelOfNormal), rejected in common germline filtering step (CommonGermline) or rejected in allele count filtering step (AlleleCount). The lower panels show scatterplots of VAF of the variants in the tumor vs VAF in the normal, VAF in the normal vs depth (DP) at the position in the normal, VAF in the tumor vs depth in the tumor.

**Supplemental Figure S12:**
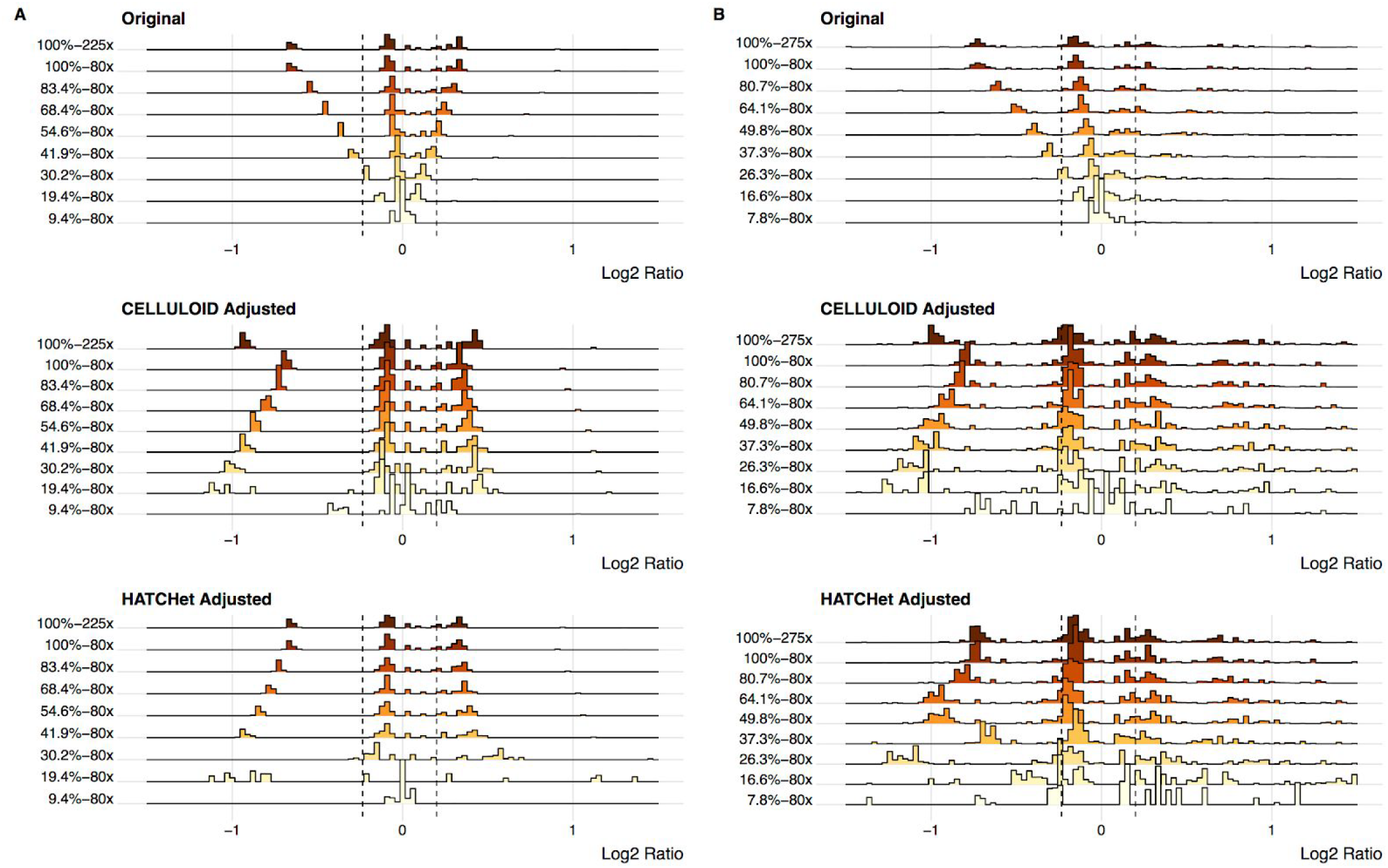
Adjustment of Log2 Values in Cell Line Purity Ladder Density plot showing the log2 values of CNVs called in the purity ladder cell lines for (A) COLO-829 and (B) HCC-1143. The first row shows the original unadjusted log2 values that were called at various purities. The second row shows the CELLULOID adjusted log2 values at the same purity levels. The third row shows the HATCHet adjusted log2 values at the same purity levels.

**Supplemental Figure S13:**
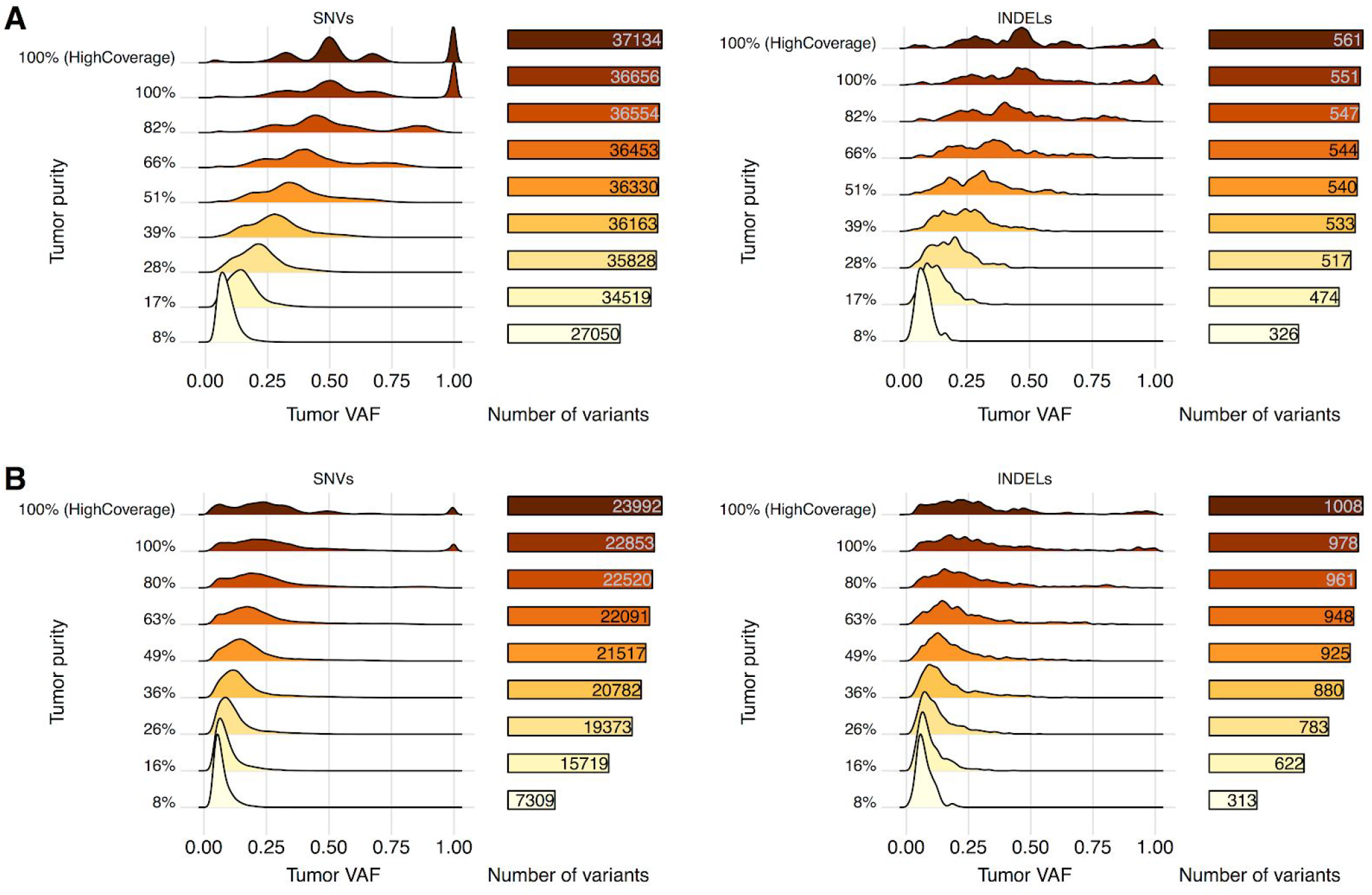
Variant allele frequency distribution and number of high confidence SNVs and Indels called in the high coverage data that are also called in the AllSomatic callsets of the purity ladder samples for (A) COLO-829 and (B) HCC-1143.

**Supplemental Figure S14:**
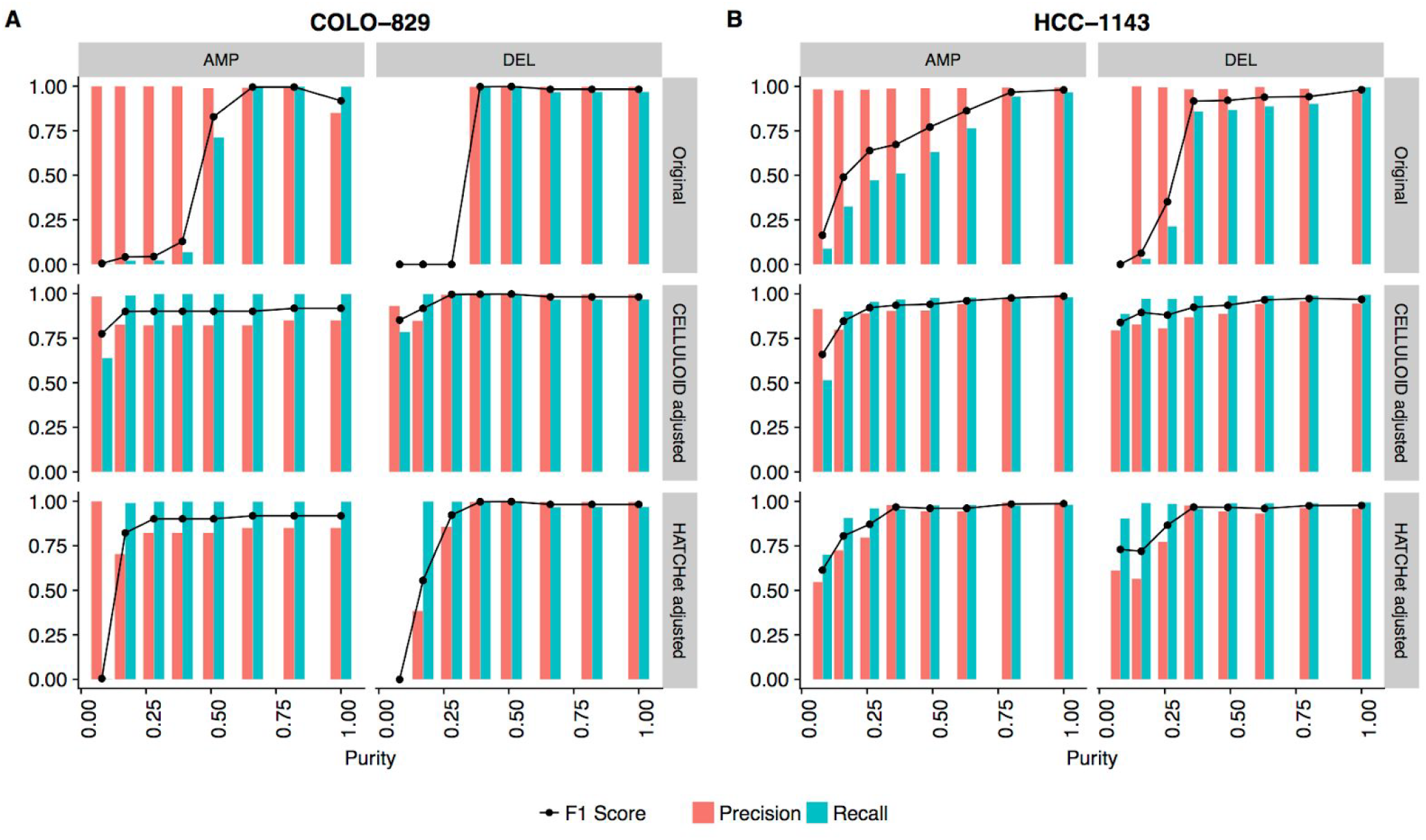
Precision, recall and F1 scores at different simulated purities for CNVs without (Original) and with (CELLULOID/HATCHet) adjustments of log2 values for purity and ploidy. Panel A corresponds to COLO-829, Panel B to HCC-1143.

**Supplemental Figure S15:**
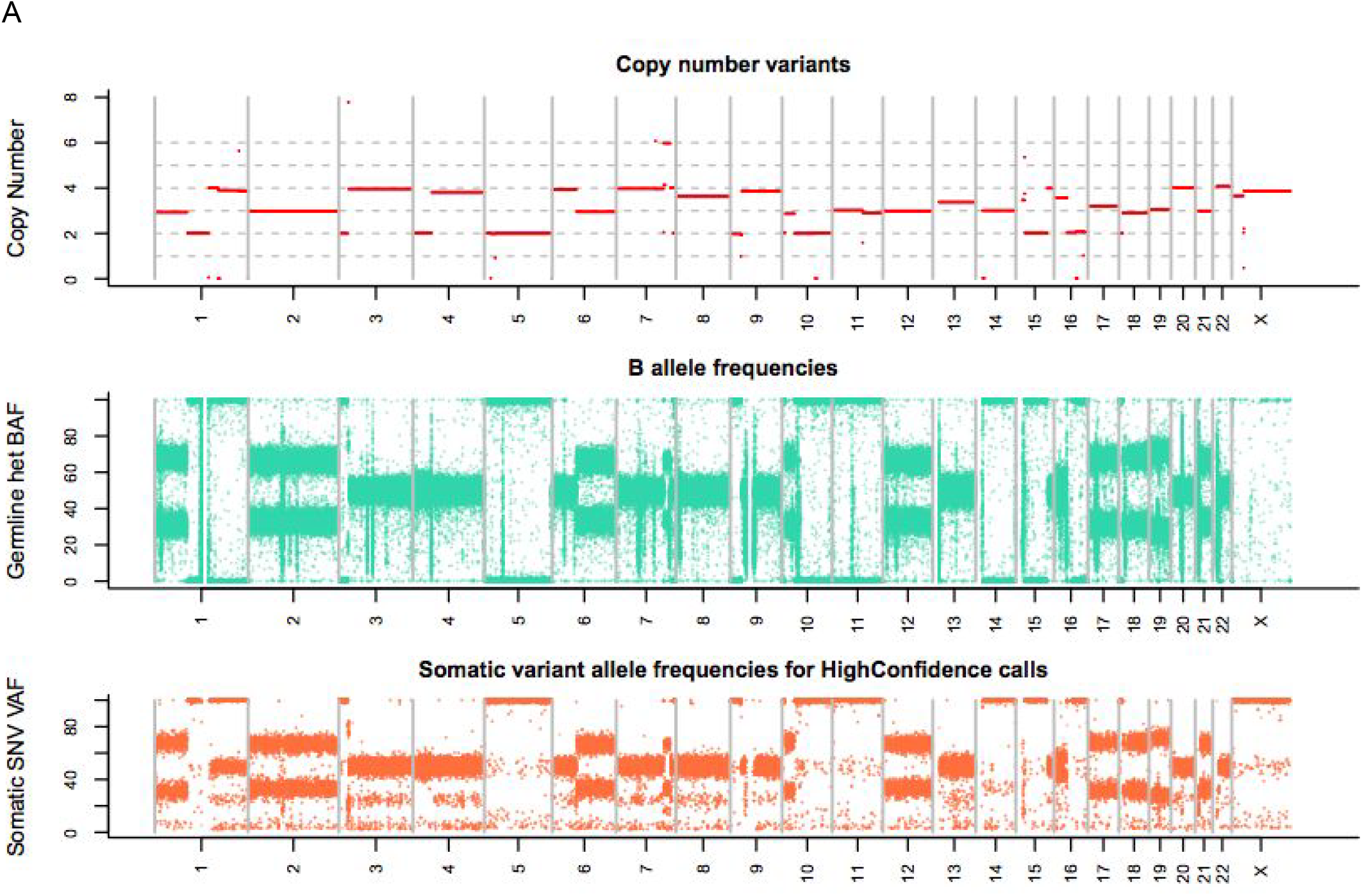
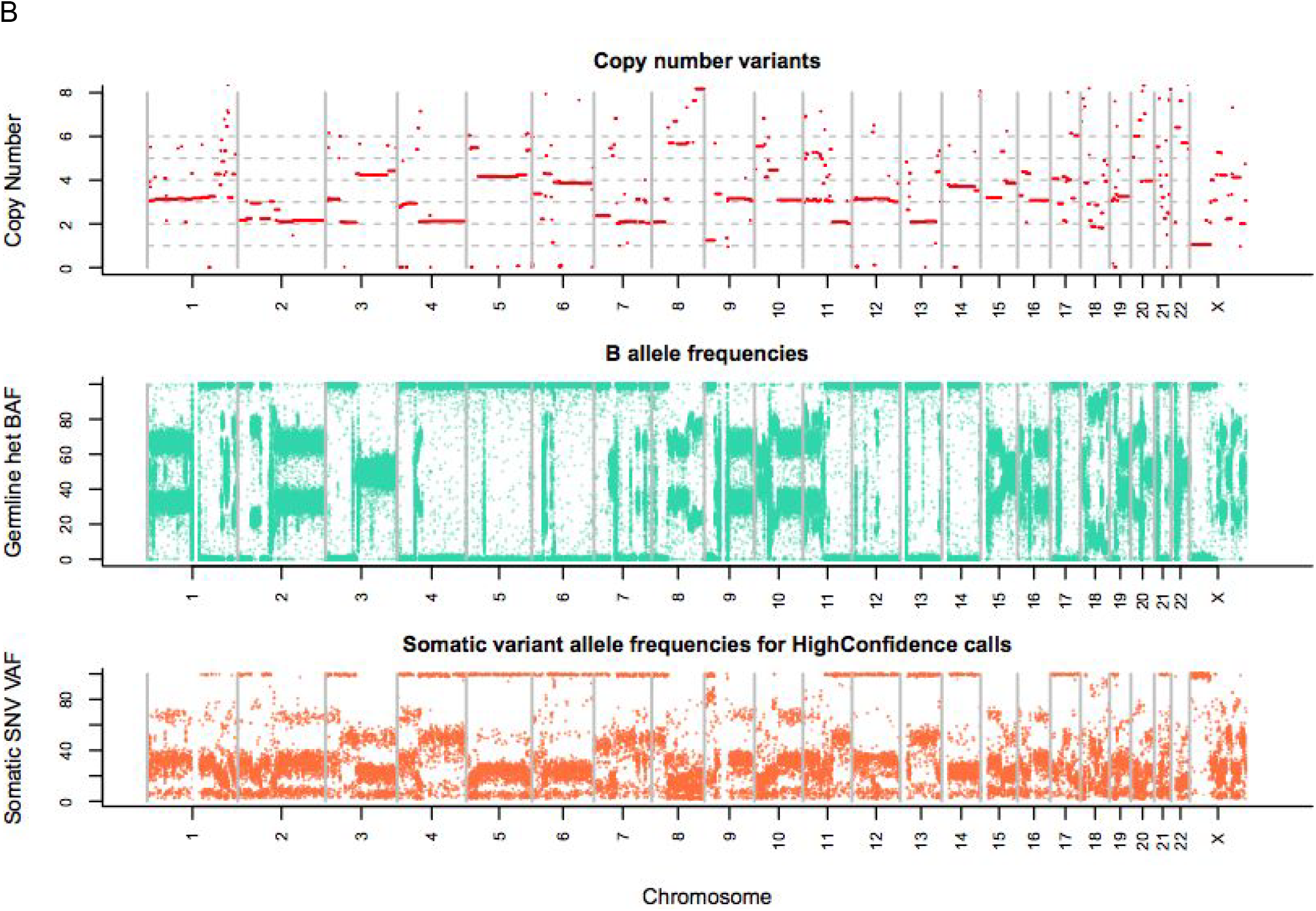
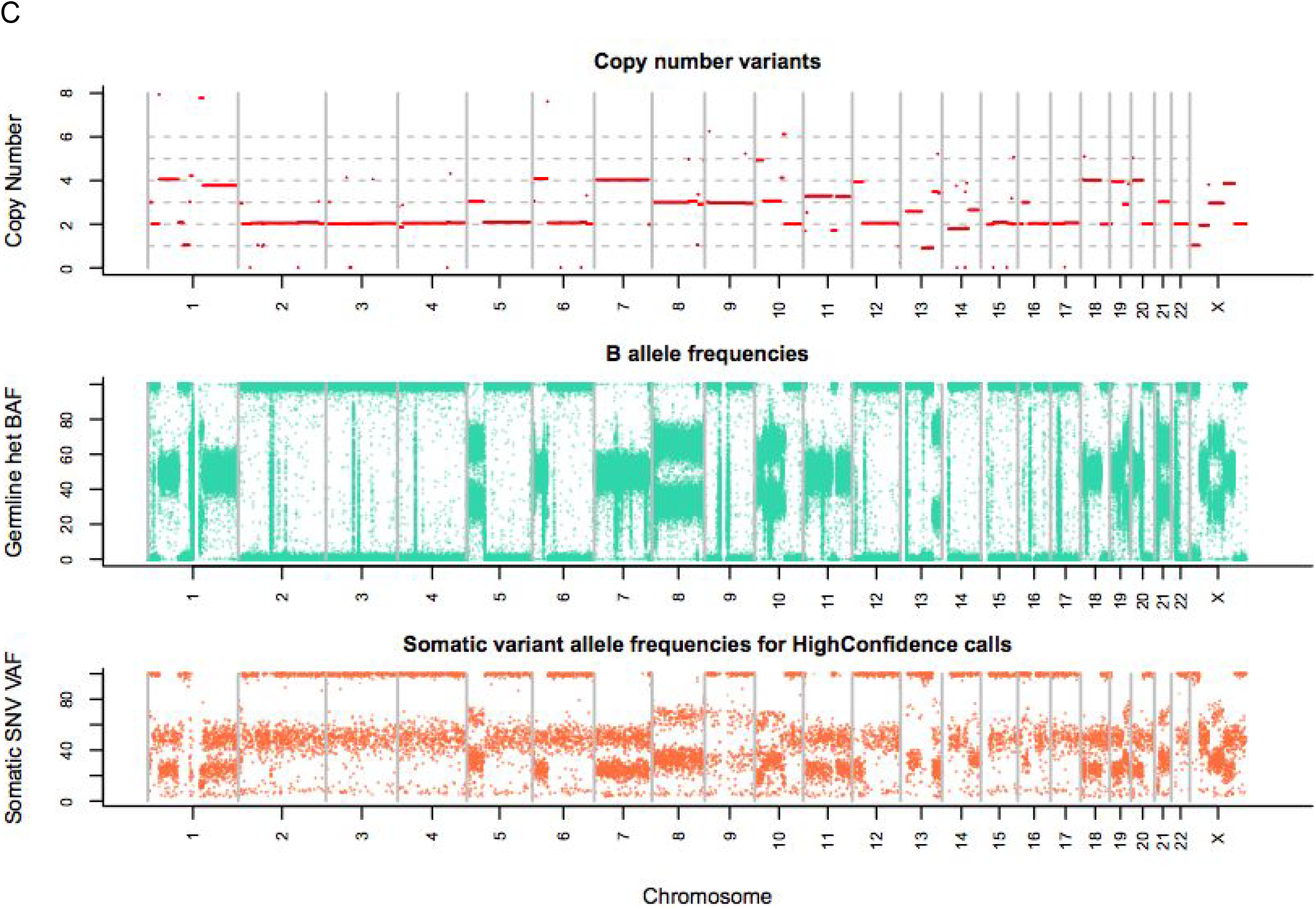
Copy number profile (top), B-allele frequency (BAF) of germline heterozygous SNPs by position (middle) and variant allele frequency (VAF) by position of HighConfidence somatic SNVs called on the high coverage NovaSeq data for (A) COLO-829, (B) HCC-1143 and (C) HCC-1187.

2 https://www.atcc.org

3 https://support.illumina.com/content/dam/illumina-support/documents/documentation/chemistry_documentation/samplepreps_truseq/truseq-dna-pcr-free-workflow/truseq-dna-pcr-free-workflow-reference-1000000039279-00.pdf

4 available at https://github.com/nygenome/nygc-short-alignment-marking

5 https://github.com/Illumina/Polaris

6 https://dockstore.org/containers/quay.io/pancancer/pcawg-sanger-cgp-workflow:develop

7 https://github.com/raphael-group/hatchet commit 0e626b0

8 https://www.nygenome.org/bioinformatics/3-cancer-cell-lines-on-2-sequencers/

